# Iron deficiency modulates metabolic landscape of Bacteroidetes promoting its resilience during inflammation

**DOI:** 10.1101/2022.11.11.516241

**Authors:** Janina P. Lewis, Qin Gui

**Affiliations:** Philips Institute for Oral Health Research; Department of Microbiology and Immunology; Department of Biochemistry, Virginia Commonwealth University, Richmond, VA, USA

## Abstract

Bacteria have to persist in low iron conditions in order to adapt to host’s nutritional immunity. Since the knowledge of iron stimulon of Bacteroidetes is sparse, we examined oral (*Porphyromonas gingivalis* and *Prevotella intermedia*) and gut (*Bacteroides thataiotaomicron*) representatives for their ability to adapt to iron deplete and iron replete conditions. Our transcriptomics and comparative genomics analysis shows that many iron-regulated mechanisms are conserved within the phylum. Those include genes upregulated in low iron: *fldA* (flavodoxin)*, hmu* (hemin uptake operon) and loci encoding ABC transporters. Downregulated were: *frd* (ferredoxin), *rbr* (rubrerythrin)*, sdh* (succinate dehydrogenase/fumarate reductase), *vor* (oxoglutarate oxidoreductase/dehydrogenase), and *pfor* (pyruvate:ferredoxin/ flavodoxin oxidoreductase). Some genus-specific mechanisms, such as the *B. thetaiotaomicron*’s *sus* coding for carbohydrate metabolism and the *xusABC* coding for xenosiderophore utilization, were also identified. While all bacteria tested in our study had the *nrfAH* operon coding for nitrite reduction and were able to reduce nitrite levels present in culture media, the expression of the operon was iron dependent only in *B. thetaiotaomicron.* It is noteworthy that we identified a significant overlap between regulated genes found in our study and the *B. thetaiotaomicron* colitis study (Zhu et al; Cell Host Microbe 27: 376-388). Many of those commonly regulated genes were also iron regulated in the oral bacterial genera. Overall, this work points to iron being the master regulator enabling bacterial persistence in the host and paves the way for more generalized investigation of the molecular mechanisms of iron homeostasis in Bacteroidetes.

## Introduction

Iron is emerging as a critical component of the microbiome-liver fat axis ^1, 2^. Furthermore, iron plays a role in development of periodontitis through it’s contribution to selection of anaerobic pathogenic bacteria in the periodontal pocket ^3, 4^. The above conditions are associated with increased levels of bacteria belonging to the Bacteroidetes phylum. The Bacteroidetes phylum are one of the four most abundant phyla in both, the oral cavity ^5, 6, 7^ and the gut ^8, 9, 10, 11^. The main characteristic is that those bacteria can grow under microaerophilic/anaerobic conditions and they rely on iron-based metabolism for energy generation ^12^. Indeed, the central enzyme, pyruvate ferredoxin oxidoreductase (PFOR) ^13^ required for decarboxylation of pyruvate to acetyl-CoA contains four iron-sulfur clusters peripherally located on the surface of the enzyme that are easily oxidized and deactivated under atmospheric oxygen levels ^13^. That group of bacteria also contains many other enzymes mediating metabolic functions that are critically dependent on iron. Thus, it is intuitive that iron levels would play a central role in regulation of metabolism of the bacteria.

Thus far, we have focused our efforts of determination of iron homeostasis on the oral bacterium *Porphyromonas gingivalis*. This is an oral pathogen shown to play a role in development and progression of periodontal disease. We have identified the major hemin uptake locus, *hmu*^14^ as well as reported on the iron stimulon of the bacterium ^15^.

We also expanded our studies onto the related oral bacterium *Prevotella intermedia* ^16^ where we found similar genes/proteins as in *P. gingivalis* that were deregulated depending on iron levels. However, no comprehensive approaches were undertaken to investigate changes in gene regulation in response to iron levels in the latter. Furthermore, the iron-dependent transcriptome of the intestinal Bacteroidetes is yet to be determined. So far, the outer membrane proteins of *B. fragilis* and *B. vulgatus* were investigated for differential expression, however no comprehensive approach to determine iron-dependent differential gene expression has been reported^17^. Finally, the molecular mechanisms as well as how the iron-dependent regulation in oral Bacteroidetes compares to iron-dependent regulation in related bacteria residing in the gut is yet to be determined.

Here we used RNAseq followed by comparative genomics to determine the iron stimulon in three Bacteroidetes species: *B. thetaiotaomicron, P. gingivalis* and *P. intermedia.* We demonstrate that there is a significant overlap in adaptation to the iron-rich and iron-deficient conditions as evidenced by common set of genes regulated in all the bacteria studied. Unique mechanisms were also detected and mainly were confined to the *B. thetaiotaomicron* bacterium.

## Materials and Methods

### Bacterial strains and growth conditions

For our study, we have used three bacterial species: *Porphyromonas gingivalis* (strains W83 and ATCC33277), *Prevotella intermedia* (strain OMA14), and *Bacteroides thetaiotaomicron* (strain BT5482 Δ*tdk*).

Two of the species are oral bacteria while the *B. thetaiotaomicron* is a gut bacterium. All bacteria are clinically significant and the *P. intermedia* OMA14 is a clinical isolate that has recently been sequenced^18^. We also include a laboratory strain of *P. gingivalis,* the ATCC33277 strain for comparison with the more virulent strain W83. The bacteria were maintained on TSB agar plates (40 g/L tryptic soy agar base supplemented with 10 g/L of yeast extract, 1g/L cysteine, 5 µg/mL hemin, and 5 µg/mL menadione). Broth cultures were prepared using enriched BHI medium (37 g/L BHI, 2.5 g/L yeast extract, 5 µg/mL hemin, and 0.5 µg/mL menadione). Iron-deplete conditions were generated by supplementing BHI with 150 µM of dipyridyl (DP). Cells were harvested in mid log phases (at an OD_660_ of approx. 0.5).

### RNA isolation and library preparation

The RNA isolation and library preparation were done as described previously ^19, 20^. Briefly, the RNeasy mini-kit (Qiagen) was used to isolate RNA from bacterial pellets following manufacturer’s protocol. The RNA was treated with the DNA-free DNase kit (Ambion,Thermo Fisher) following manufacturer’s protocol to remove any remaining DNA. The quality of RNA was verified using agarose gel electrophoresis to ensure that the RNA was intact and then used to generate RNA-seq library using the Ovation Complete Prokaryotic RNA-Seq DR multiplex kit (Nugen, currently Tecan) following the manufacturer’s instruction. The libraries were submitted to the VCU’s Nucleic Acid Sequencing core facility where the quality of the libraries was verified using the bioanalyzer (Agilent Technologies).

### Highthroughput sequencing and data analysis

Libraries that were verified and had acceptable quality were sequenced by the VCU’s Nucleic Acid Sequencing core using the MiSeq Illumina Genome Analyzer (*P. gingivalis* and *P. intermedia* libraries) and with the Illumina HiSeq 4000 (*B. thetaiotaomicron*). The sequencing data were then analyzed with the CLC Genomic Workbench V12 (CLC Bio, Qiagen). Thus, resulting transcriptome-derived sequences were aligned to the reference genomes of: *P. gingivalis* W83 (for *P. gingivalis* W83 transcriptome), *P. gingivalis* ATCC33277 (for *P. gingivalis* ATCC33277 transcriptome), *P. intermedia* OMA14 (for the *P. intermedia* OMA14 transcriptome) and to *B. thetaiotaomicron* VPI BT5482 (for the *B. thetaiotaomicron* VPI BT5482 Δ*tdk* transcriptome). The enrichment for the iron-dependent regulation was determined by dividing the number of reads derived from the bacteria grown in iron-deplete conditions to that derived from the bacteria grown under iron replete conditions for each position of the genome following normalization by aligned fragment per kilobase of transcript per million (FPKM)^21, 20, 19^.

### Bioinformatics analysis

The BioCyc genome viewer was used to align genomic loci for selected genes ^22^. The alignment of similar loci was done across the three different bacterial species used in our study. The reference genomes used were: *P. gingivalis* W83, *P. intermedia* 17 (the *P. intermedia* OMA14 genome is not available on BioCyc) and the *B. thataiotaomicron* BT5482.

### Nitrite reduction assay

Bacterial cultures were grown in Tryptic Soy Broth (TSB) or BHI supplemented with 0.5 mg/ml cysteine, 5 µg/mL hemin, and 0.5 µg/mL menadione. Iron-deplete conditions were generated by supplementing the media with 150 µM of dipyridyl (DP). Nitrite was added at 0.3 and 2 mM respectively. Bacterial growth was monitored by measuring optical density at 660nm and nitrite levels were determined using the Griess reagent according to manufacturer’s instruction (Abcam). Cultures were initiated by inoculating BHI media with bacteria prepared on fresh blood agar plates. Actively growing, overnight cultures were then diluted 1:30 (for *P. gingivalis* W83 and *P. intermedia* OMA14) or 1:50 (for *B. thetaiotaomicron*) and grown anaerobically for 72hr, at 37°C. 1ml aliquots were removed at different time points and used to determine bacterial growth and levels of nitrite. As a control, media with and without nitrite were maintained in parallel with the bacterial cultures.

### Statistical analysis

All experiments were repeated on at least four separate days (biological replicates). For the analysis of the changes at the levels of mRNA under iron replete and iron deplete conditions the fold change and P values were derived from the four biological replicates. T test was used to determine statistical significance for the nitrite assays.

## Results

### Iron-dependent stimulon of P. gingivalis

Genes that were the most upregulated in *P. gingivalis* W83 grown under iron-depleted conditions as compared to iron replete conditions are listed in Table 1. 96 genes were upregulated 1.4 fold or more. Among those genes 11 operons were identified. The most highly upregulated gene was PG0827 coding for a MATE family efflux transporter (920 fold upregulation). This gene has been reported to also have cis-encoded antisense transcript indicating that it may be regulated by small RNA ^23^. The PG0495 coding for a T9SS type A sorting domain-containing protein was upregulated 87 fold. Also, several transporter-encoding operons were upregulated and included: the PG1551-6 operon encoding the Hmu hemin uptake system (upregulated 11.4 - 30.1 fold), PG1175 – PG1181 coding for a two component lipoprotein/MMPL transporter followed by ABC transporter ATP-binding proteins as well as the TetR/AcrR family transcriptional regulator (upregulated 13.4 – 136.2 fold), PG1019-1022 TonB-dependent transporter system (upregulated by 20.8 – 214.5 fold, depending on a gene). It is noteworthy that genes coding for proteins mediating metabolic processes were also regulated. The most highly upregulated in this class was the PG1857-58 operon coding for proteins with unknown function (PG1857 - 48 fold) and flavodoxin (PG1858 - 133 fold). Also, the locus carrying PG1461 -1469 and encoding hypothetical proteins and putative isoprenylcysteine, carboxymethyltransferase family proteins as well as class I SAM – dependent methyltransferase were upregulated by 2 – 27 fold, depending on a gene. Genes encoding proteins that associate with iron such as PG1236-38 operon coding for hemerythrin domain containing protein: (RluA family pseudouridine synthase) and a response regulator/transcription factor was upregulated by 2-7 fold, depending on a gene. Slightly upregulated were genes coding for iron - (FeoB2, PG1294) and manganease - (FeoB1, PG1043) transport as well as the *ahpC* gene coding for peroxiredoxin (2.4 fold). Finally, genes coding for the T9SS type A sorting domain-containing protein, PG0654 and PG1374, were upregulated 1.5 and 7.3 fold, respectively.

**Table 1.**
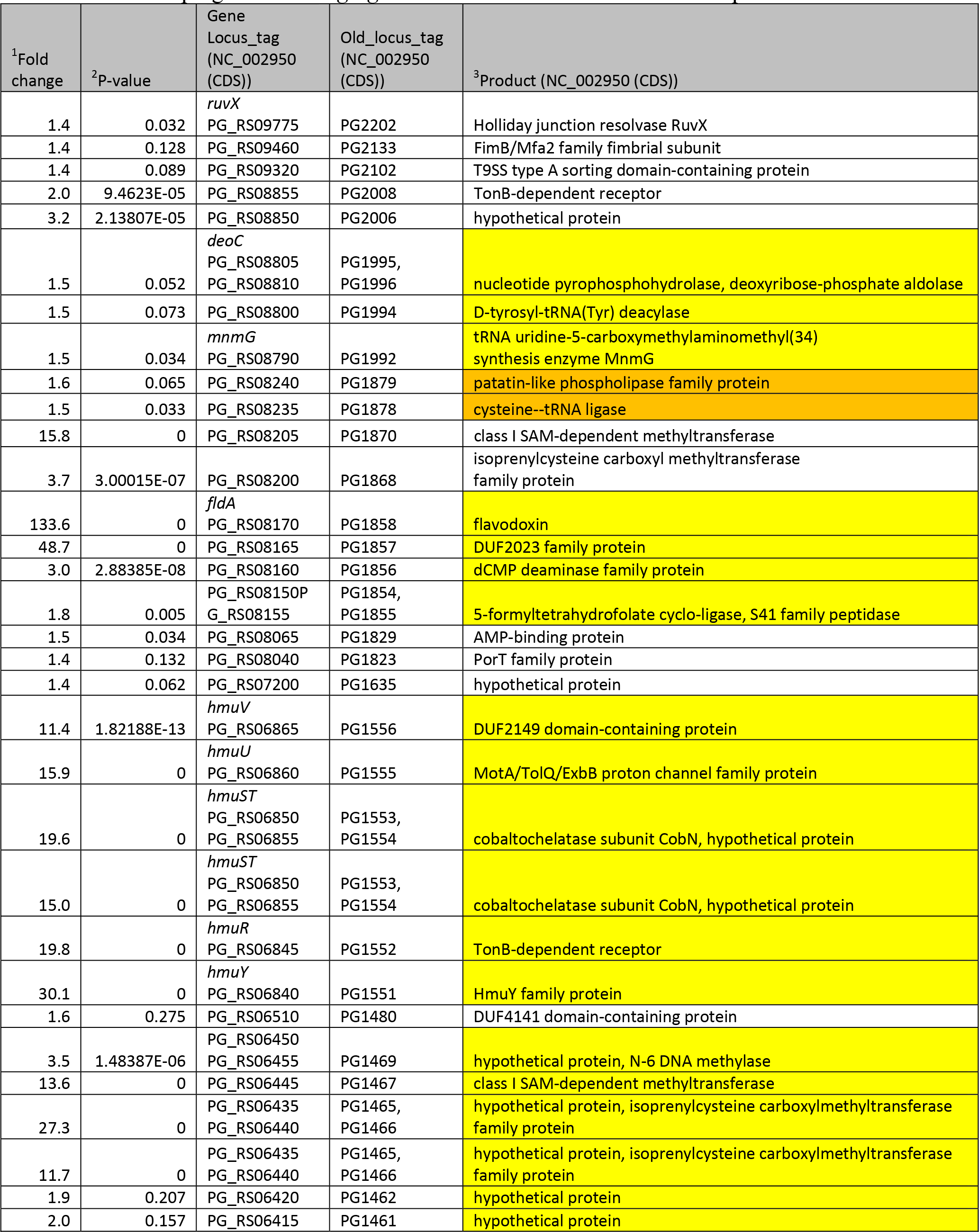
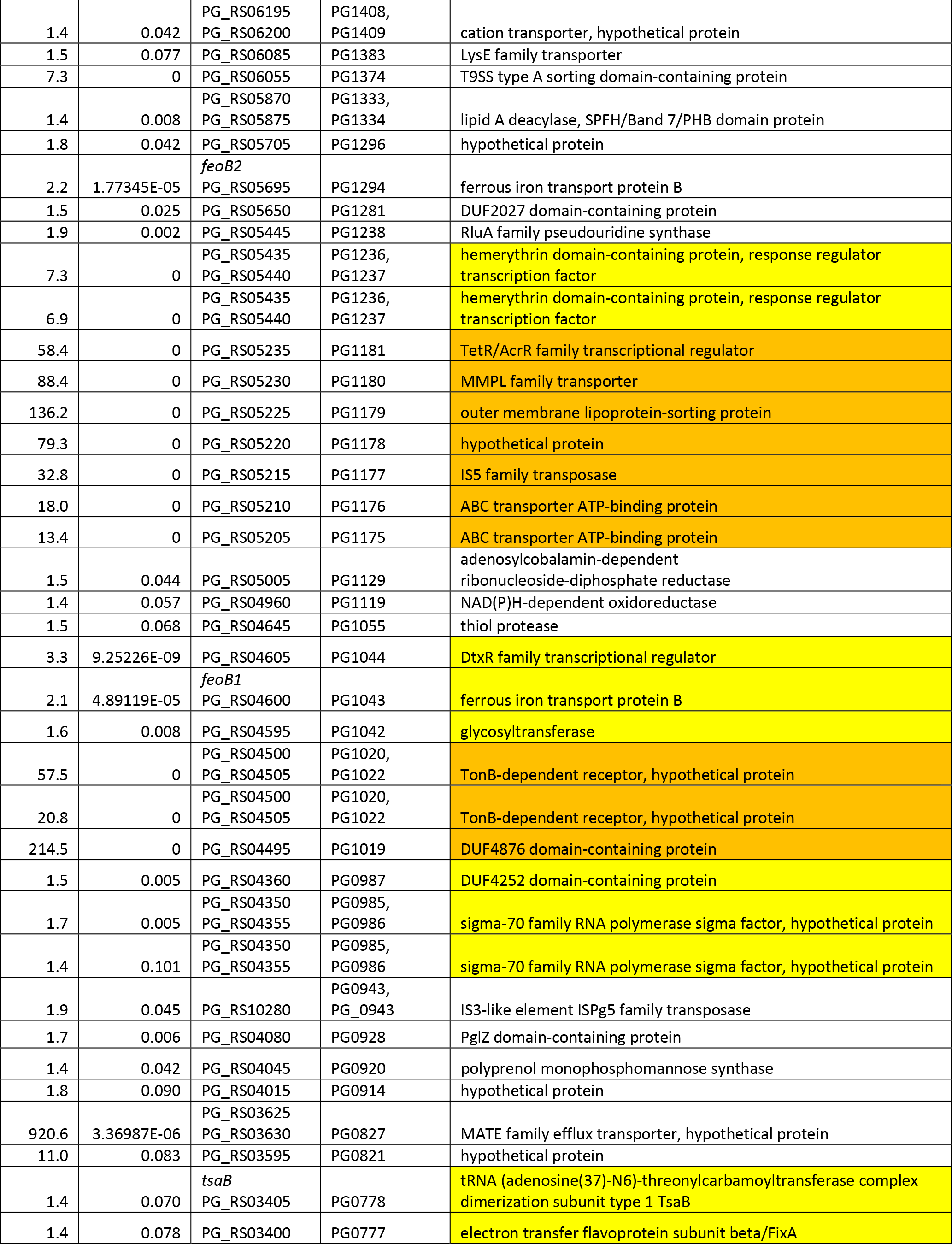
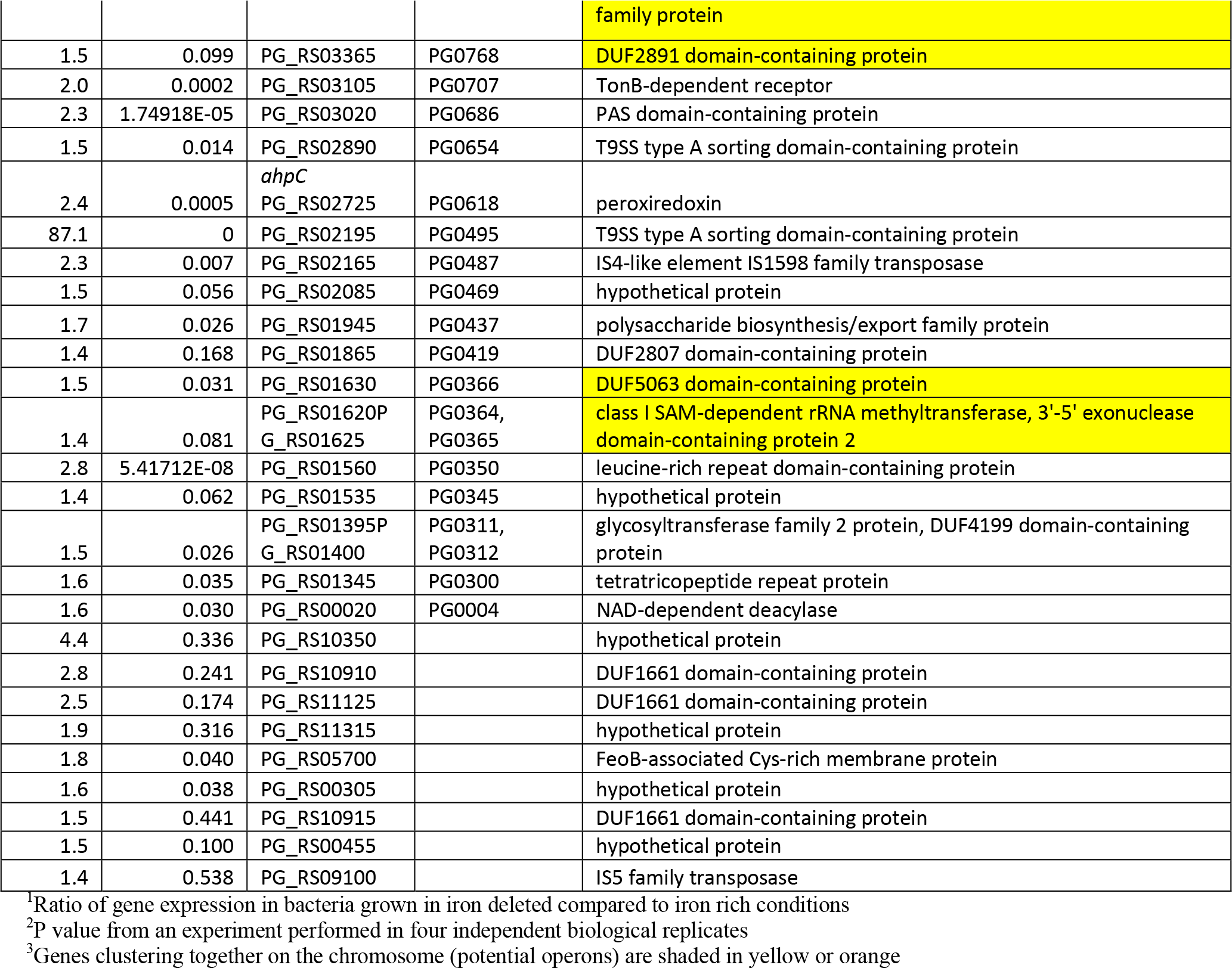
Genes upregulated in P. gingivalis W83 under iron deplete conditions (at least 1.4 fold).

Genes that were most downregulated in *P. gingivalis* W83 grown under iron-depleted conditions as compared to iron replete conditions are listed in Table 2. 67 genes were downregulated 2 fold or more (Table 2). The most highly downregulated gene, PG0195, codes for rubrerythrin and is downregulated 46 fold. Other downregulated genes encoded iron-based metabolism proteins. The first is an operon PG0302-0309 coding for an electron transport Rnf complex. It is downregulated 2-3.8 fold, depending on a gene. Also, operon PG1614-1616 coding for the fumarate reductase/succinate dehydrogenase was downregulated by 3.5 – 3.7 fold, depending on a gene. In addition, the operon PG1809-1813 coding for oxoglutarate oxidoreductase/dehydrogenase system, Vor, and flanked by a 4Fe-4S binding protein (ferredoxin) was downregulated by 5.2 – 7.4 fold, depending on a gene. Other, strikingly downregulated genes encode a hypothetical proteins (PG_RS11335, PG_RS11350, PG_RS10005 – nomenclature according to new locus tag) (30.8 fld, 9.2 fld, and 6.3 fld, respectively). No old locus tags were available for those loci (Table 2).

**Table 2.** Genes downregulated in P. gingivalis W83 under iron deplete conditions (at least 2 fold).

| <sup>1</sup>Fold change | <sup>2</sup>P-value | Gene Locus_tag (NC_002950 (CDS)) | Old_locus_tag (NC_002950 (CDS)) | <sup>3</sup>product (NC_002950 (CDS)) |
| --- | --- | --- | --- | --- |
| -30.8 | 0.058 | PG_RS11335 |  | hypothetical protein |
| -9.2 | 0.226 | PG_RS11350, PG_RS10690 |  | DUF1661 domain-containing protein |
| -6.3 | 0.055 | PG_RS10005 |  | hypothetical protein |
| -5.6 | 4.9036E-12 | PG_RS10650 |  | hypothetical protein |
| -3.8 | 0.082 | PG_RS10940, PG_RS11290 |  | DUF1661 domain-containing protein |
| -3.7 | 0.491 | PG_RS11360 |  | hypothetical protein |
| -3.1 | 0.00203383 | PG_RS10460, PG_RS06160 |  | DUF1661 domain-containing protein, transposase |
| -3.0 | 0.106 | PG_RS11285 |  | hypothetical protein |
| -2.9 | 0.045 | PG_RS10395 |  | hypothetical protein |
| -2.6 | 0.227 | PG_RS11115 |  | DUF1661 domain-containing protein |
| -2.6 | 0.338 | PG_RS10535 |  | DUF1661 domain-containing protein |
| -2.6 | 0.0003 | PG_RS04140 |  | IS5 family transposase |
| -2.4 | 0.252 | PG_RS10550 |  | DUF1661 domain-containing protein |
| -2.3 | 0.669 | PG_RS10870, PG_RS10875 |  | DUF1661 domain-containing protein, hypothetical protein |
| -2.2 | 0.302 | PG_RS11000 |  | DUF1661 domain-containing protein |
| -2.1 | 0.473 | PG_RS10580 |  | DUF1661 domain-containing protein |
| -2.1 | 0.005 | PG_RS02100 |  | DUF4248 domain-containing protein |
| -2.0 | 0.035 | PG_RS05410 |  | IS5 family transposase |
| -2.0 | 0.115 | PG_RS10200 |  | DUF1661 domain-containing protein |
| -1.9 | 0.152 | PG_RS11050 |  | IS3 family transposase |
| -1.9 | 0.212 | PG_RS10250 |  | glycine cleavage system H protein |
| -2.0 | 0.0009 | PG_RS00510 | PG0110 | glycosyltransferase |
| -2.5 | 0.135 | PG_RS00810 | PG0177 | IS4-like element IS1598 family transposase |
| -46.1 | 0 | PG_RS00900 | PG0195 | rubrerythrin family protein |
| -2.2 | 0.118 | PG_RS01030 | PG0225 | IS4-like element IS1598 family transposase |
| -2.8 | 1.3105E-07 | PG_RS01350 | PG0302 | SoxR reducing system RseC family protein |
| -3.4 | 8.9495E-13 | PG_RS01355 | PG0303 | Fe-S cluster domain-containing protein |
| -3.4 | 3.8814E-08 | rsxC PG_RS01360 | PG0304 | electron transport complex subunit RsxC |
| -3.8 | 2.7878E-12 | PG_RS01365, PG_RS01370 | PG0305, PG0306 | RnfABCDGE type electron transport complex subunit D, RnfABCDGE type electron transport complex subunit G |
| -3.6 | 1.5993E-10 | PG_RS01365, PG_RS01370, PG_RS01375 | PG0305, PG0306, PG0307 | RnfABCDGE type electron transport complex subunit D, RnfABCDGE type electron transport complex subunit G, electron transport complex subunit E |
| -3.1 | 3.19E-09 | PG_RS01370, PG_RS01375 | PG0306, PG0307 | RnfABCDGE type electron transport complex subunit G, electron transport complex subunit E |
| -2.0 | 0.008 | PG_RS01380 | PG0308 | electron transport complex protein RnfA |
| -2.3 | 1.8935E-05 | PG_RS01385, PG_RS01390 | PG0309, PG0310 | FAD:protein FMN transferase, nitroreductase family protein |
| -2.0 | 0.127 | PG_RS11275 | PG0442, PG_0442 | hypothetical protein |
| -4.1 | 0.195 | PG_RS02045 | PG0460 | IS5-like element ISPg8 family transposase |
| -3.6 | 0.042 | PG_RS10105 | PG0524, PG_0524 | DUF1661 domain-containing protein |
| -2.2 | 0.077 | PG_RS10115, PG_RS10120 | PG0545, PG_0545, PG0546, PG_0546 | restriction endonuclease subunit S, helix-turn-helix domain-containing protein |
| -2.1 | 0.064 | PG_RS02500, PG_RS02505 | PG0563, PG0564 | hypothetical protein |
| -4.3 | 4.4923E-10 | PG_RS04540 | PG1031 | IS5 family transposase |
| -2.0 | 1.5145E-08 | PG_RS05185,<br>PG_RS05190 | PG1171, PG1172 | (Fe-S)-binding protein, lactate utilization protein |
| -2.0 | 0.0002 | PG_RS05185,<br>PG_RS05190 | PG1171, PG1172 | (Fe-S)-binding protein, lactate utilization protein |
| -2.6 | 0.044 | PG_RS05275 | PG1197 | IS5 family transposase |
| -1.9 | 0.0007 | *nrdG* PG_RS05545 | PG1259 | anaerobic ribonucleoside-triphosphate reductase activating protein |
| -2.0 | 0.139 | PG_RS06340 | PG1445 | RteC domain-containing protein |
| -3.1 | 4.5733E-06 | PG_RS06360 | PG1448 | IS5 family transposase |
| -2.1 | 0.186 | PG_RS06425 | PG1463 | hypothetical protein |
| -2.0 | 0.002 | PG_RS06610 | PG1500 | helicase |
| -2.1 | 0.196 | PG_RS10525 | PG1527,PG_1527 | helix-turn-helix domain-containing protein |
| -2.0 | 0.005 | PG_RS06820 | PG1545 | superoxide dismutase |
| -1.9 | 0.056 | PG_RS06925 | PG1573 | Crp/Fnr family transcriptional regulator |
| -3.6 | 2.5498E-12 | *sdhB*, PG_RS07115 | PG1614 | succinate dehydrogenase/fumarate reductase iron-sulfur subunit |
| -3.7 | 5.5302E-12 | *sdhA*, PG_RS07120 | PG1615 | fumarate reductase/succinate dehydrogenase flavoprotein subunit |
| -3.5 | 5.6732E-14 | *sdhC*, PG_RS07125 | PG1616 | succinate dehydrogenase/fumarate reductase cytochrome b subunit |
| -5.2 | 0 | PG_RS07975 | PG1809 | 2-oxoacid:acceptor oxidoreductase family protein |
| -5.1 | 1.3323E-15 | PG_RS07980 | PG1810 | 2-oxoglutarate oxidoreductase |
| -5.6 | 1.8874E-15 | *vorB* PG_RS07985 | PG1812 | 3-methyl-2-oxobutanoate dehydrogenase subunit VorB |
| -7.4 | 0 | PG_RS07990 | PG1813 | 4Fe-4S binding protein |
| -2.8 | 0.084 | PG_RS10710,<br>PG_RS11190 | PG1980,PG_1980 | DUF1661 domain-containing protein |
| -3.0 | 0.013 | PG_RS11390 | PG2114,PG_2114 | hypothetical protein |
| -2.0 | 0.184 | PG_RS09375 | PG2115 | DUF1661 domain-containing protein |
| -2.5 | 0.008 | PG_RS09740 | PG2194 | IS4-like element IS1598 family transposase |

<sup>1</sup>Ratio of gene expression in bacteria grown in iron deleted compared to iron rich conditions
<sup>2</sup>P value from an experiment performed in four independent biological replicates
<sup>3</sup>Genes clustering together on the chromosome (potential operons) are shaded in yellow or orange

The conclusion from the above-described regulated genes is that iron/hemin uptake and iron – independent metabolism mechanisms are drastically upregulated while the iron-dependent metabolism and oxidative stress defense mechanisms are downregulated under iron-chelated conditions. These results were also verified using the *P. gingivalis* ATCC33277 (see supplemental Tables 1 and 2). Of note were also 3 DUF1661 domain-containing protein coding genes (upregulated by 9, 3.7, 3 and >2 fold) and 13 DUF1661 domain – containing protein encoding genes were downregulated. Most of those are genes coding for small proteins specific for *P. gingivalis* (except one encoded by PG0174 [PG_RS00800]). Fig. 3E.

Iron-dependent stimulon of P. intermedia

The most drastically regulated genes in *P. intermedia* OMA14 grown without iron as compared to the bacterium grown in iron replete conditions are listed in Tables 3 and 4. There was 101 genes upregulated at least 2 fold in *P. intermedia* OMA14 (Table 3). Among the most upregulated were ones encoding metabolic proteins: PIOMA14_I_0020-21 (*fldA,* flavodoxin) and DUF2023 family protein (159 and 164 fold, respectively). Also, there were multiple operons coding for TetR/AcrR family transcriptional regulator drastically upregulated in low iron conditions. Thus, on the large chromosome significantly upregulated was operon PIOMA14_I_0603-5 coding for a TonB-dependent receptor, hypothetical protein, and a TetR/AcrR family transcriptional regulator (129-, 289-, 268- fold, respectively). Also, an operon PIOMA14_I_0908-14 coding for: TetR/AcrR family transcriptional regulator, two ABC transporter ATP-binding proteins, MMPL family transporter, outer membrane lipoprotein-sorting protein, hypothetical protein, and hypothetical protein, site-specific integrase was significantly upregulated (131-37 fold, depending on a gene in the operon). Furthermore, an operon PIOMA14_I_1565 – 1572 encoding: TetR/AcrR family transcriptional regulator, ABC transporter ATP-binding protein, ABC transporter ATP- binding protein, MMPL family transporter, outer membrane lipoprotein-sorting protein, hypothetical protein, hypothetical protein, site-specific integrase was upregulated by 40, 81, 223, 106, 76, 99, and 54 fold, respectively depending on the gene. Finally, facing in opposite direction, an operon with similar composition as regards the proteins encoded by the genes in the locus PIOMA14_I_1509 – 1516 including: TetR/AcrR family transcriptional regulator, ABC transporter ATP-binding protein, ABC transporter ATP- binding protein, MMPL family transporter, outer membrane lipoprotein-sorting protein, hypothetical protein, hypothetical protein (putative site-specific integrase) upregulated by: 43, 96, 136, 108, 48, 99, and 48 fold, respectively. Present on the small chromosome was an operon PIOMA14_II_0447-9 coding for TetR/AcrR family transcriptional regulator, ABC transporter ATP-binding protein, and ABC transporter ATP-binding protein that was upregulated by 246, 102, and 851 fold, respectively.

**Table 3.**
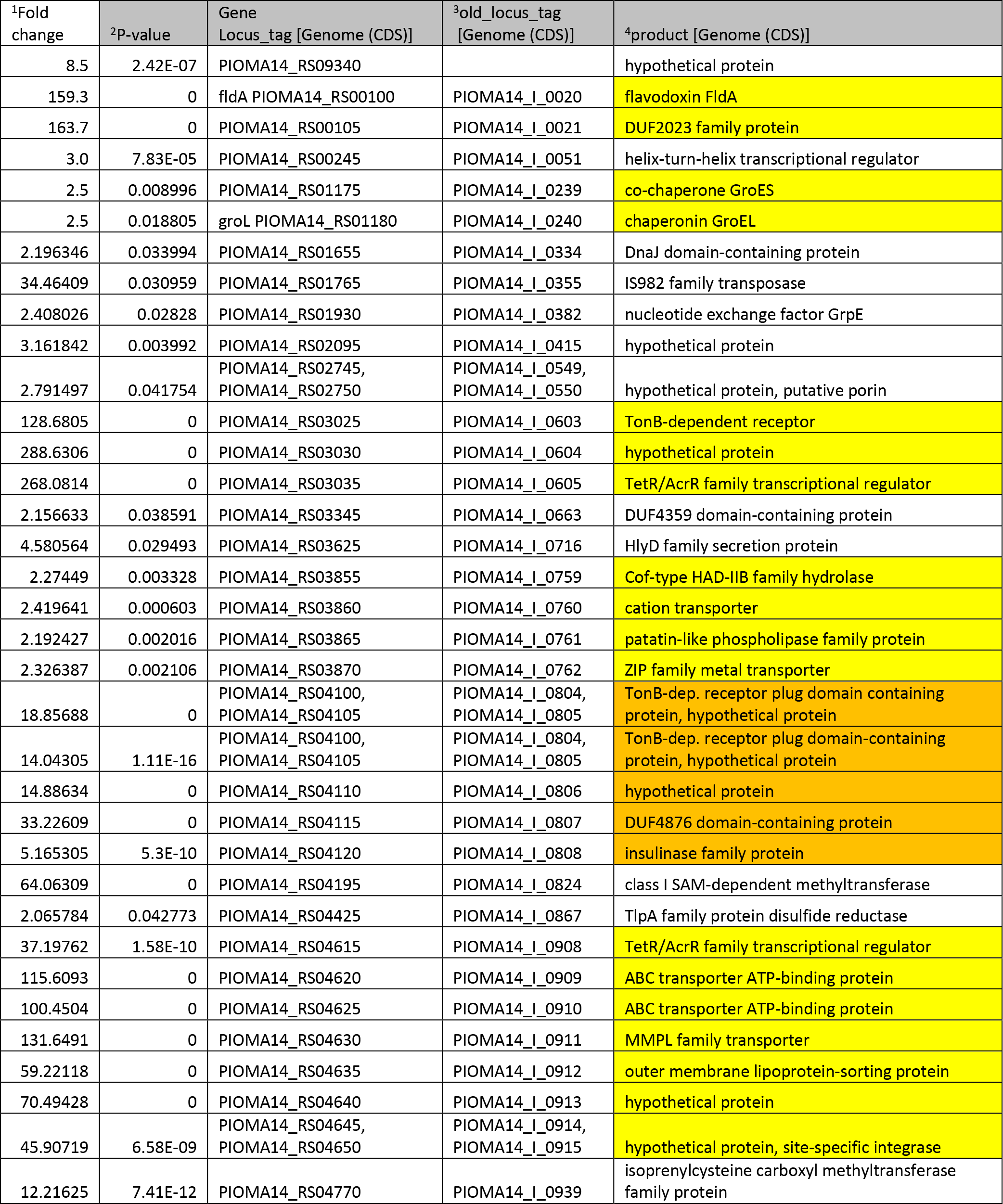
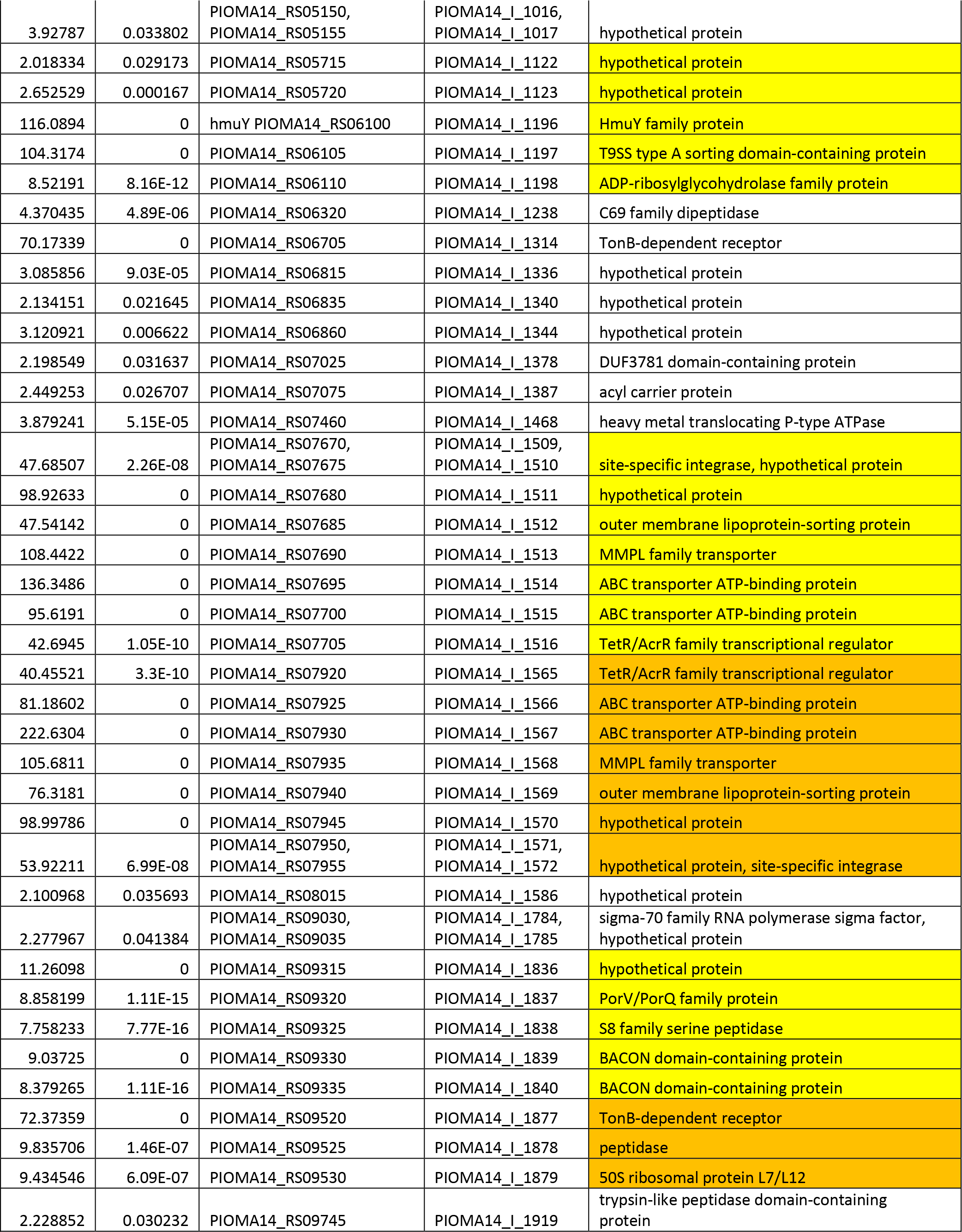
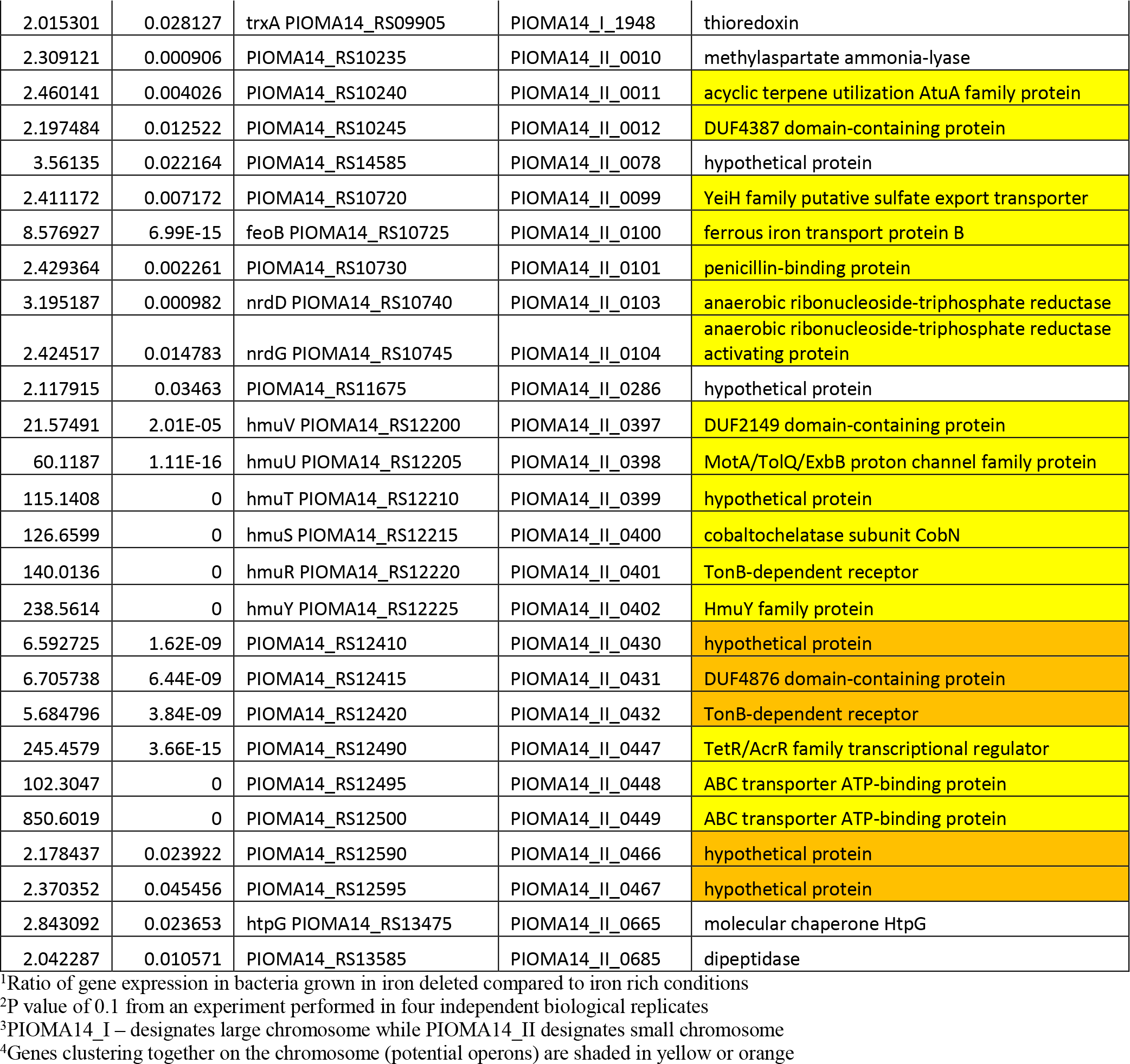
Genes upregulated in *P. intermedia* OMA14 under iron deplete conditions (at least 2 fold).

**Table 4.**
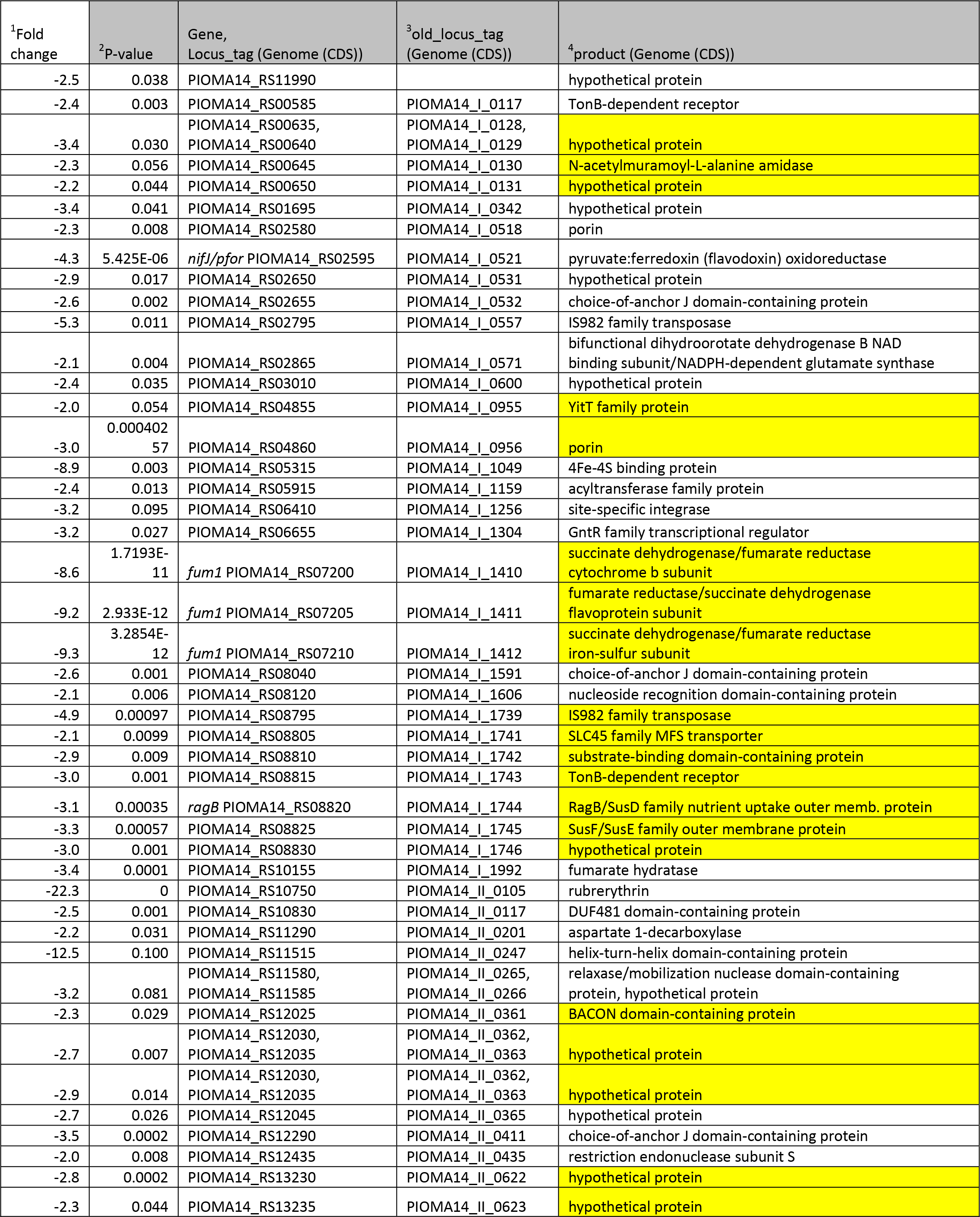
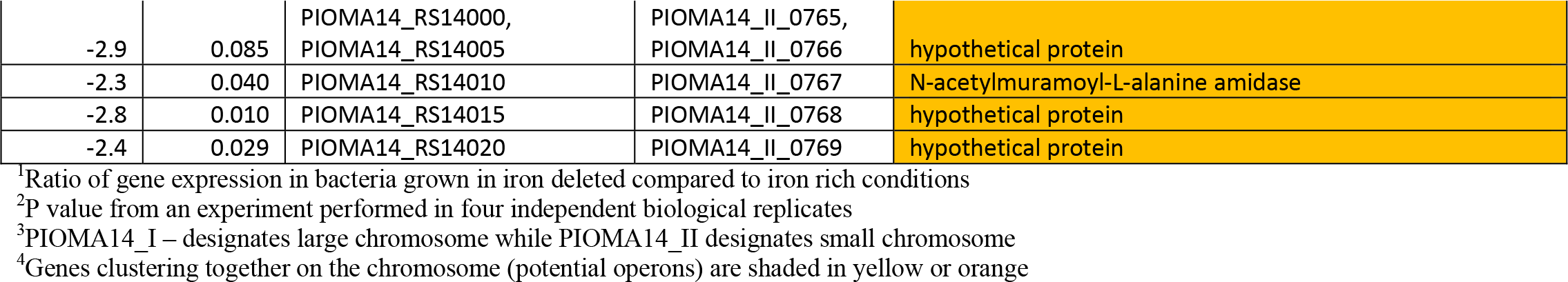
Genes downregulated in P. intermedia OMA14 under iron deplete conditions (at least 2 fold).

Two hemin uptake loci were significantly upregulated. A locus PIOMA14_I_1196 (PIOMA14_RS06100) coding for the hemin uptake receptor HmuY, followed by T9SS type A sorting domain-containing protein, and ADP-ribosylglycohydrolase family protein were upregulated 116-, 104- and 8.5 fold, respectively. A second *hmu* hemin uptake locus located on the small chromosome PIOMA14_II_0397 – 402 and encoding: HmuY family protein, TonB-dependent receptor, cobaltochelatase subunit CobN, hypothetical protein, MotA/TolQ/ExbB proton channel family protein, and DUF2149 domain-containing protein was upregulated 239, 140, 127, 115, 60 and 22 fold respectively.

The latter operon bears significant similarity with the *P. gingivalis hmu* operon identified by Lewis et al ^14^. Of interest among the upregulated genes is also locus (PIOMA14_II_0099 – PIOMA14_II_104) coding for sulfate transporter, iron transport protein (FeoB) and anaerobic ribonucleotide-triphosphate reductase (NrdD and NrdG).

Downregulated by at least 2 fold was 49 genes (Table 4). PIOMA14_II_0105 coding for rubrerythrin was the most drastically downregulated gene (22.3 fld). Also, significantly downregulated was operon PIOMA14_I_1410 – 12 encoding the fumarate reductase/succinate dehydrogenase system (8.5, 9.2 and 9.3 fld, respectively). PIOMA14_I_1049 (PIOMA14_RS05315) coding for 4Fe-4S binding protein (ferredoxin – similar to PG1421 and BT2414) was also downregulated by 8.9 fld (Fig. 2E, Supplemental Figure 1). The major iron-dependent metabolic enzyme pyruvate:ferredoxin (flavodoxin) oxidoreductase, PIOMA14_I_0521, was downregulated by 4.3 fld. Finally, PIOMA14_II_0247 coding for the helix-turn-helix domain-containing protein was downregulated by 12.5fld. Many other genes coding for transport system such as the SusD/E system (PIOMA14_I_1739 – 46), TonB-dependent system (PIOMA14_I_0117), porin (PIOMA14_I_0518 and PIOMA14_I_0956), hypothetical proteins were downregulated (Table 4).

In summary, in *P. intermedia* iron depletion lead to overexpression of iron- independent metabolic mechanisms, iron uptake mechanisms, and downregulation of iron-based metabolism and oxidative stress response mechanisms.

### Iron-dependent stimulon of B. thetaiotaomicron

Genes regulated by iron levels in *B. thetaiotaomicron* VPI BT5482 Δ*tdk* are listed in Tables 5 and 6. We identified 323 genes upregulated by at least 2 fold (Table 5). The most drastically regulated was a locus composed of three genes BT2063 - 065 (upregulated by: 489.4, 676.1, and 663 fold, depending on a gene). The locus codes for a TonB-dependent receptor and two proteins with DUF4374 domain and with PepSY domain, respectively ^2^. This locus was recently shown to be involved in xenosiderophore utilization and highly upregulated in the colitis model^2^. In addition, highly upregulated was operon BT0491- 498 encoding the Hmu – like hemin uptake system (upregulation ranging from 108.9 – 290.8 fld, depending on a gene). BT0507 - 509 coding for TetR/AcrR family transcriptional regulator and two ABC transporter ATP-binding proteins were upregulated by 51.2, 516.2, and 675.5 fold, respectively. Highly upregulated was a locus: BT2473 – 482 encoding fimbrillin-family protein, two cytochrome c biogenesis proteins CcsA, porin, thiol oxidoreductase, peptidase M75, helix-turn-helix transcriptional regulator, and three hypothetical proteins with 69.4 – 323.7 fold upregulation depending on a gene. It is also noteworthy that the BT2479 was also upregulated in the colitis model ^2^. A locus BT3625 - 633 coding for proteins with PepSY-associated TM helix domain, DUF4857 domain, fimbrillin protein, TonB-dependent receptor, ABC transporter ATP-binding protein, and hypothetical protein was upregulated by 36.7 – 128 fold, depending on a gene within the locus.

**Table 5.**
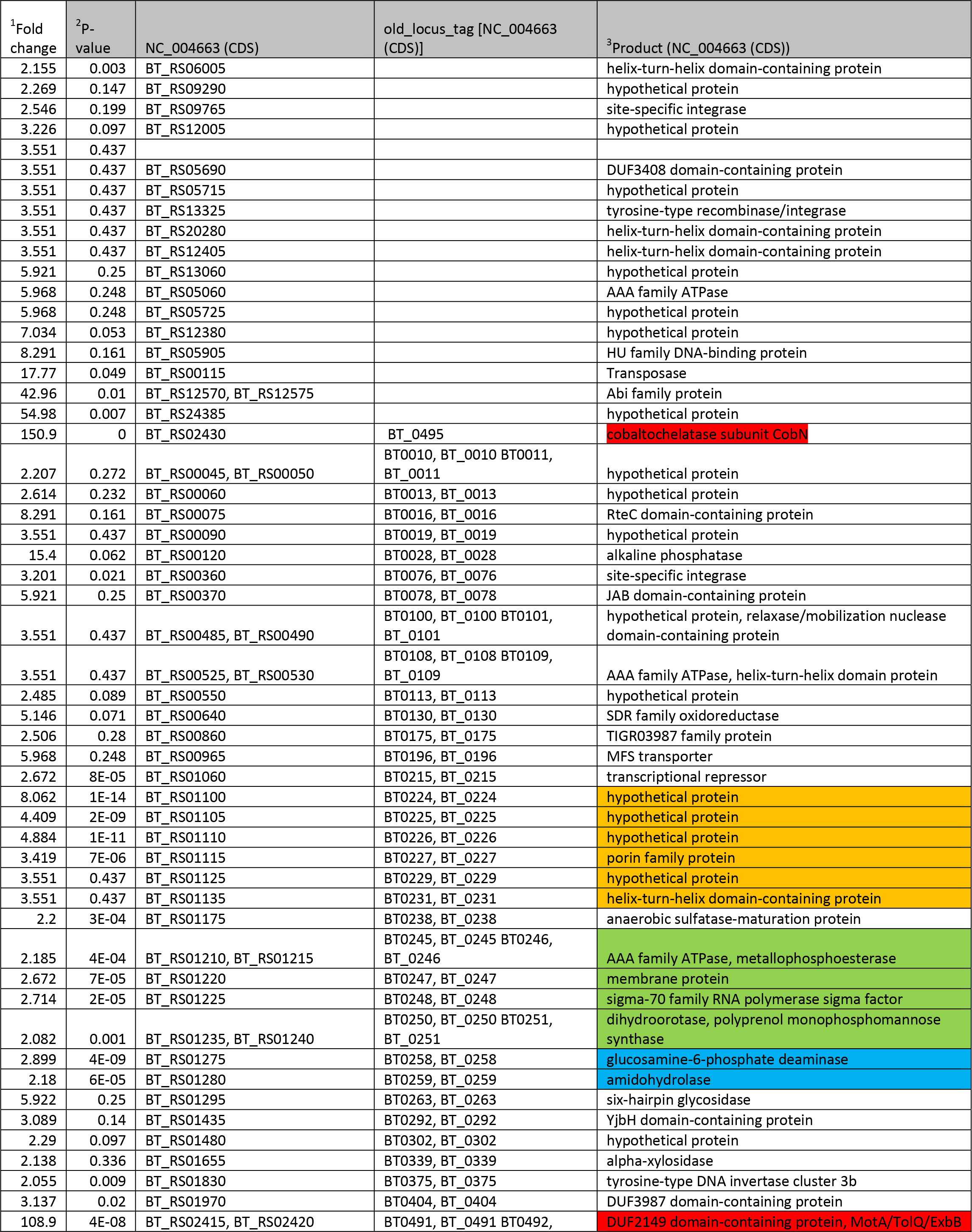
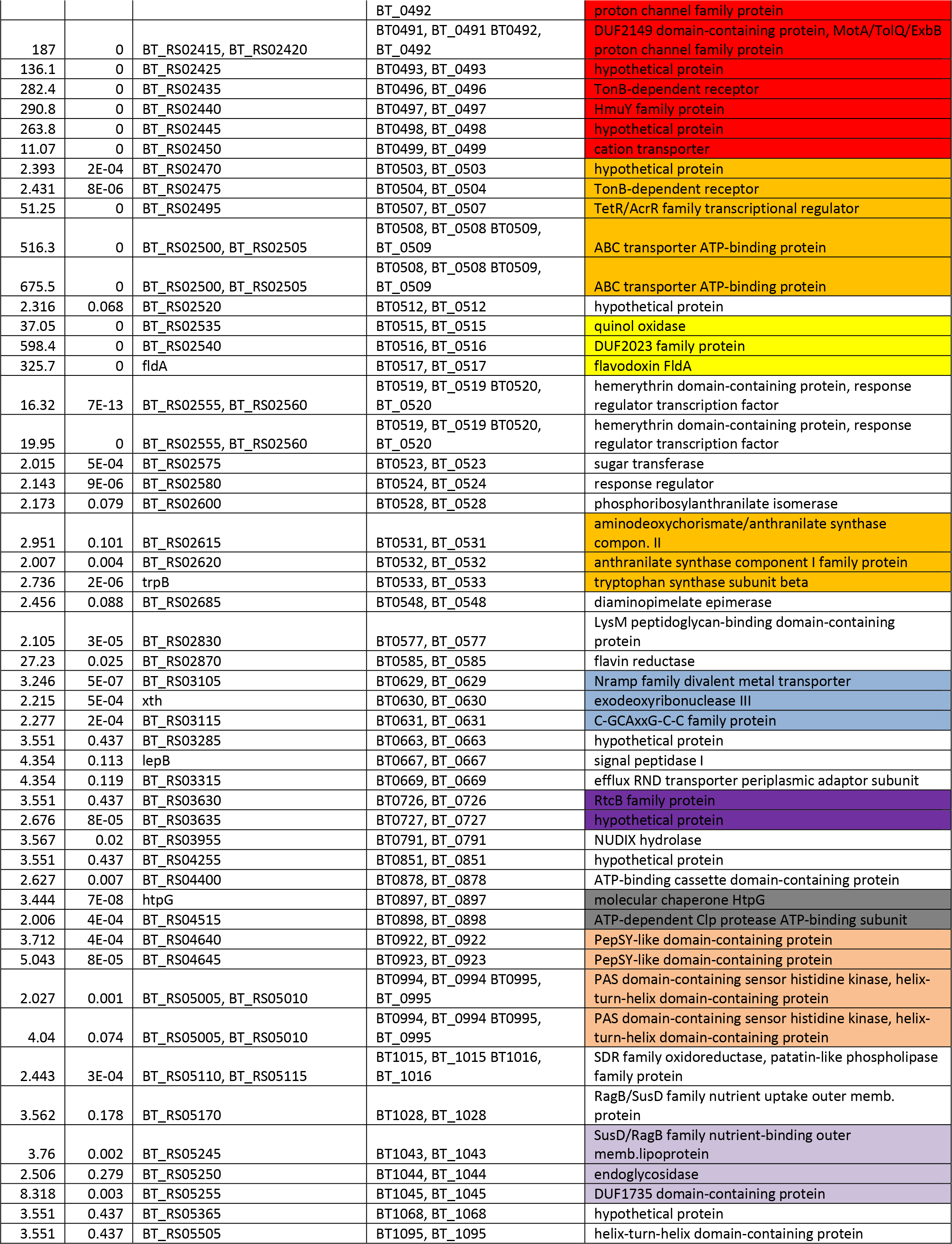
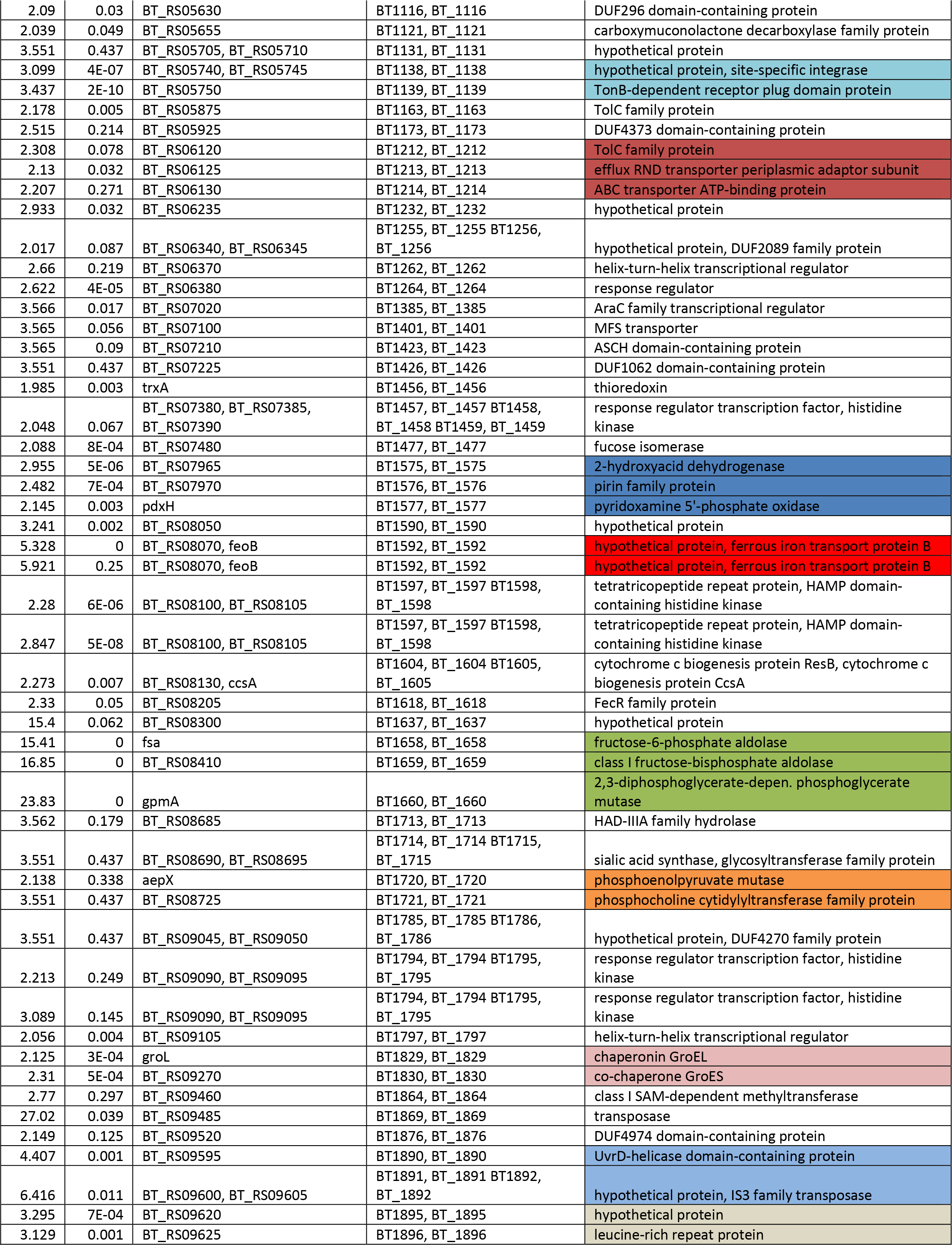
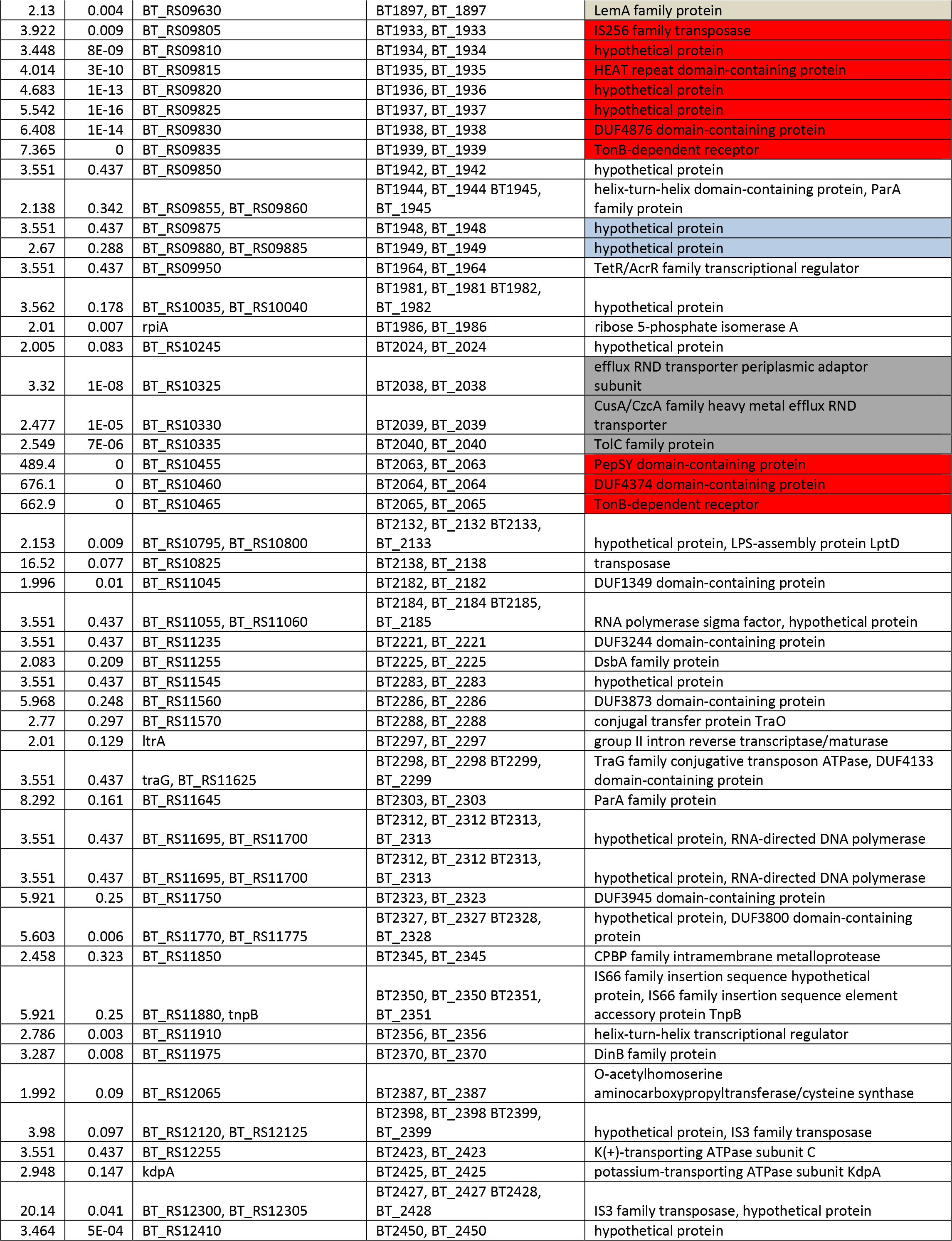
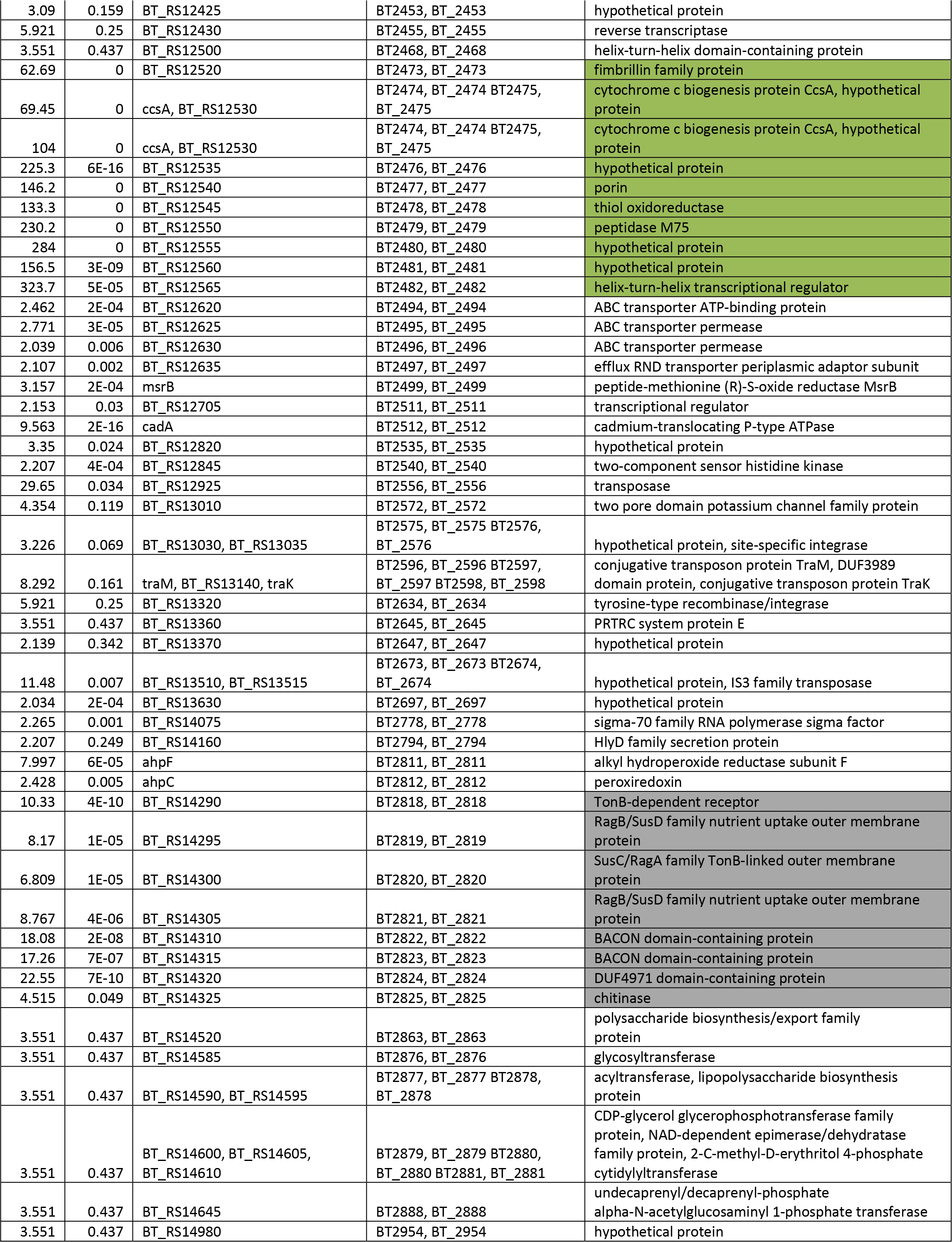
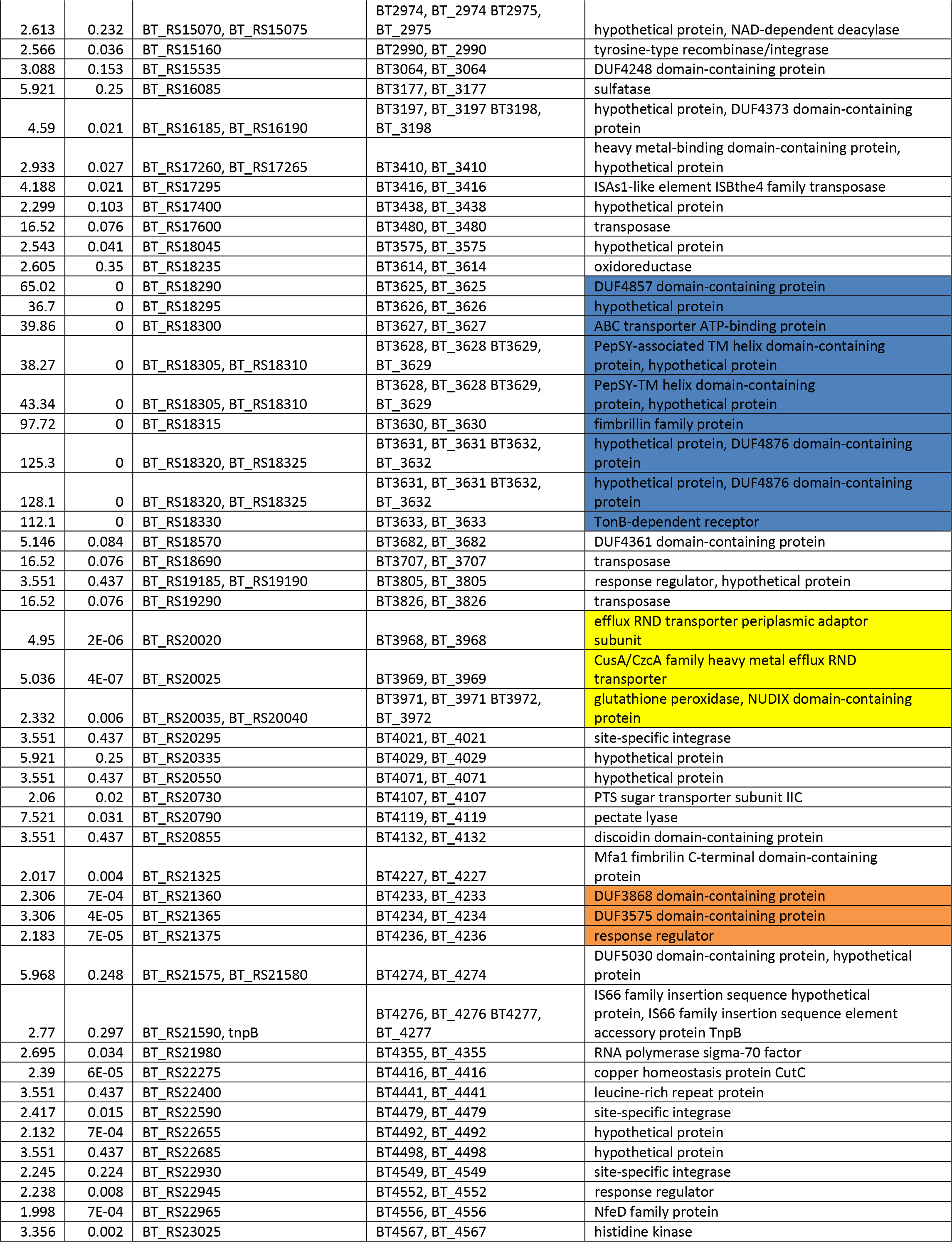
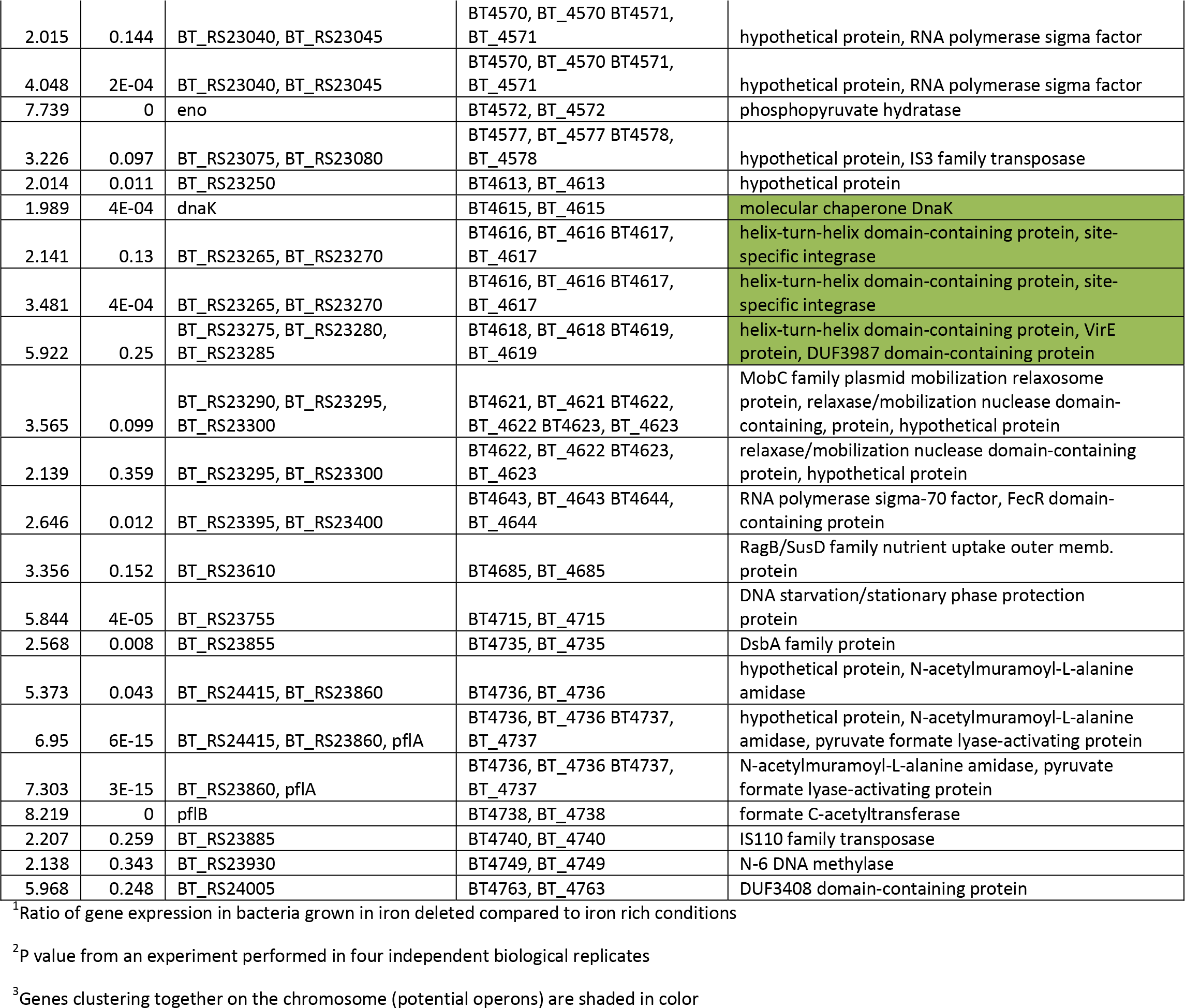
Genes upregulated in B. thetaiotaomicron BT5482 Δtdk at least 2 fold under iron deplete conditions.

**Table 6.**
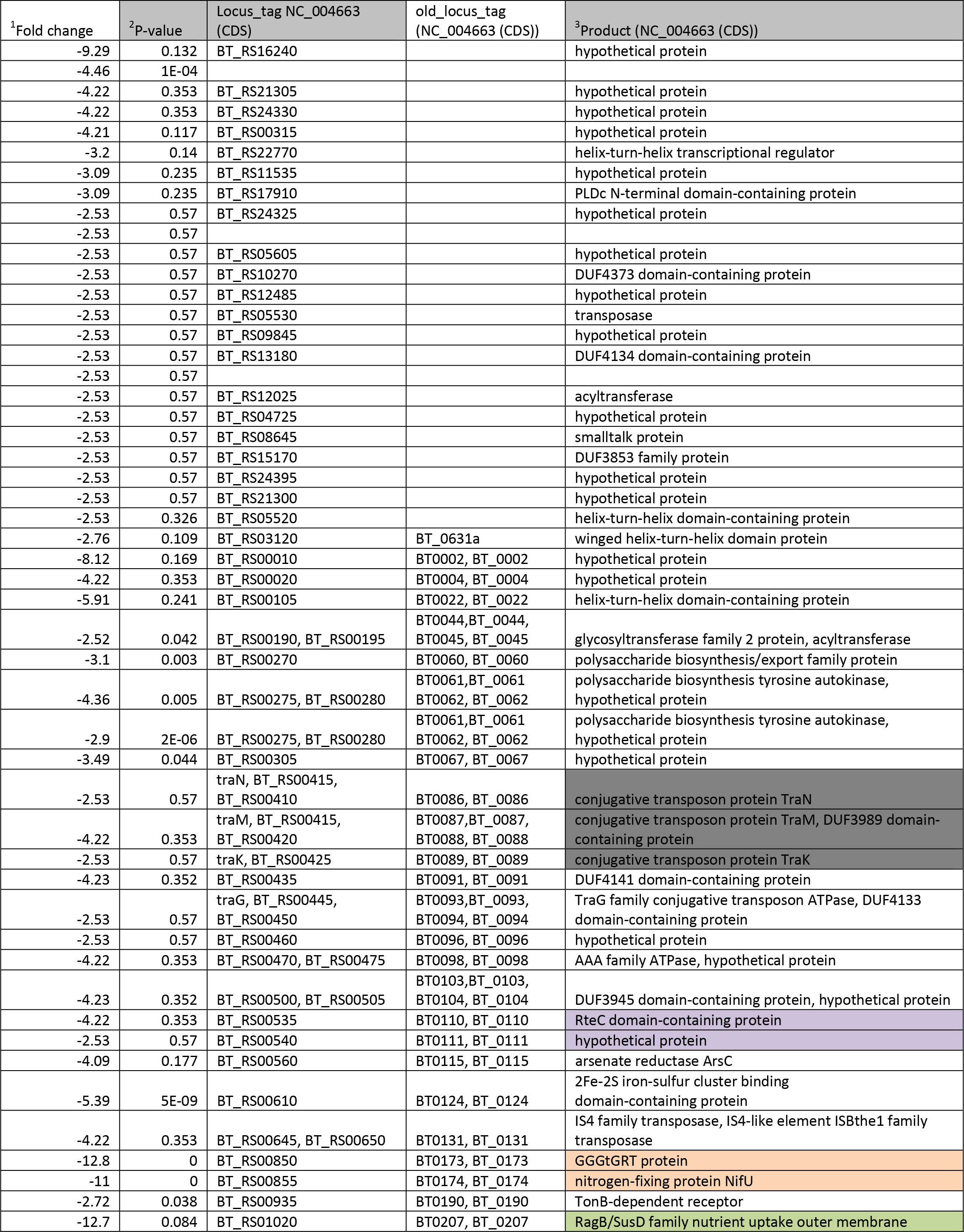
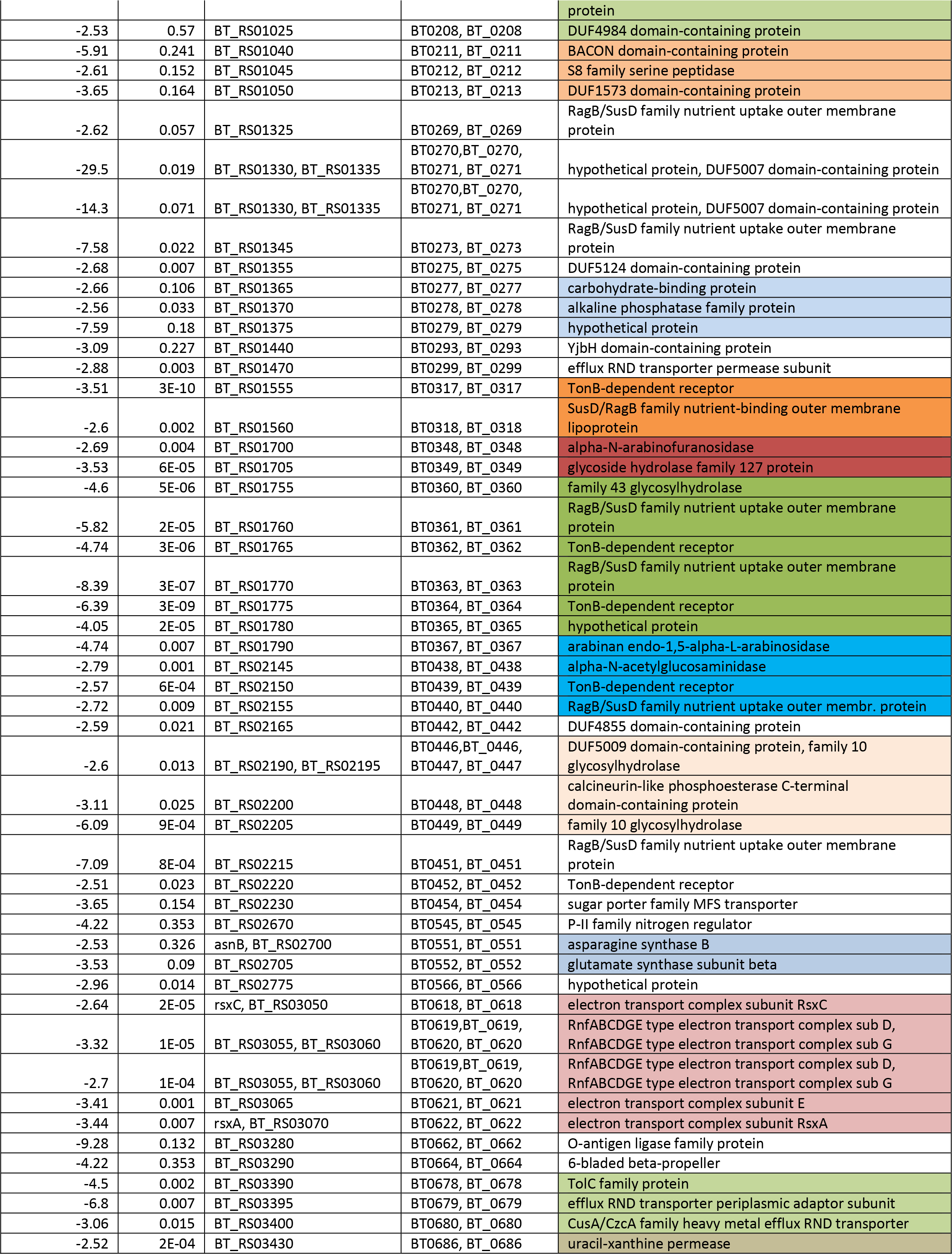
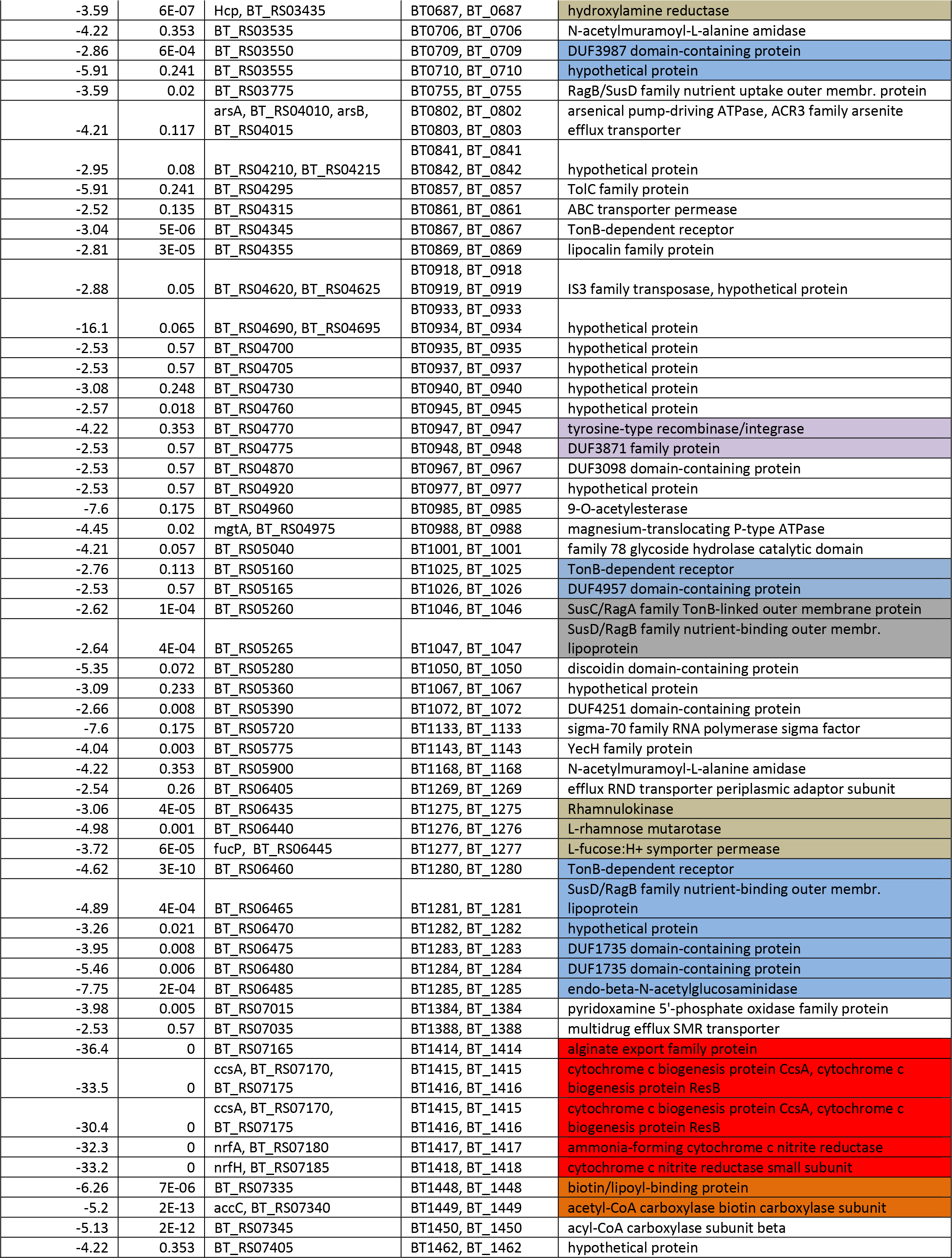
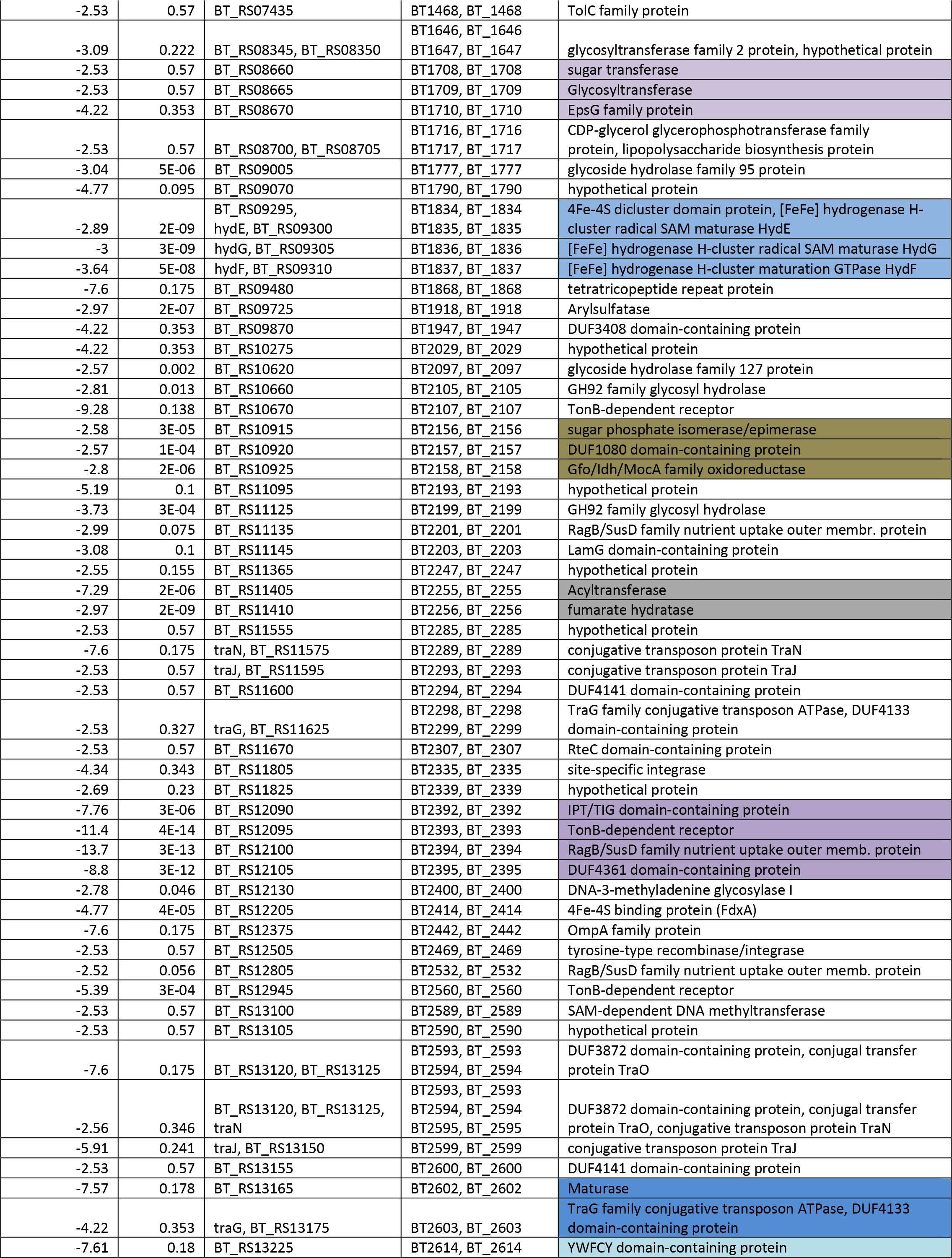
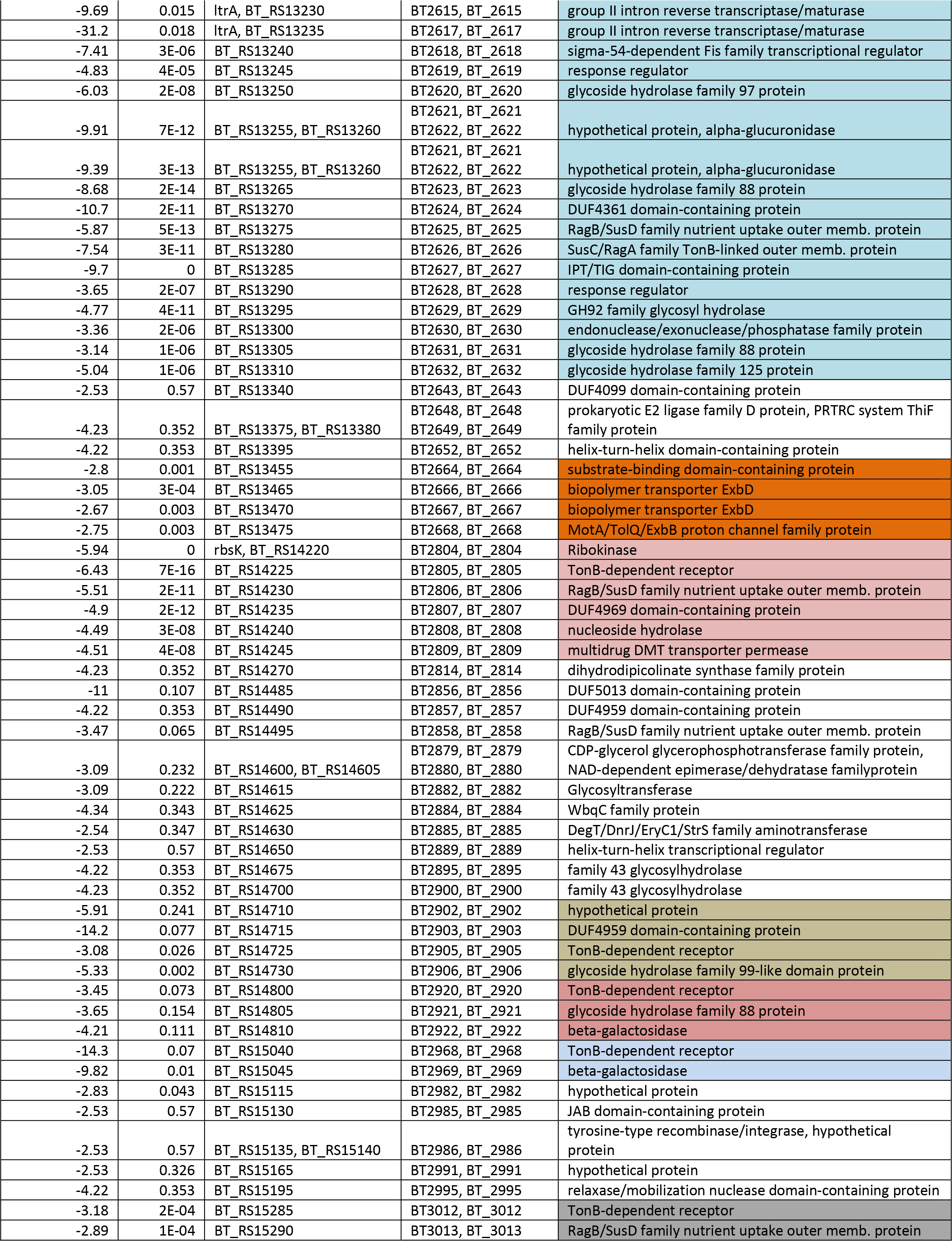
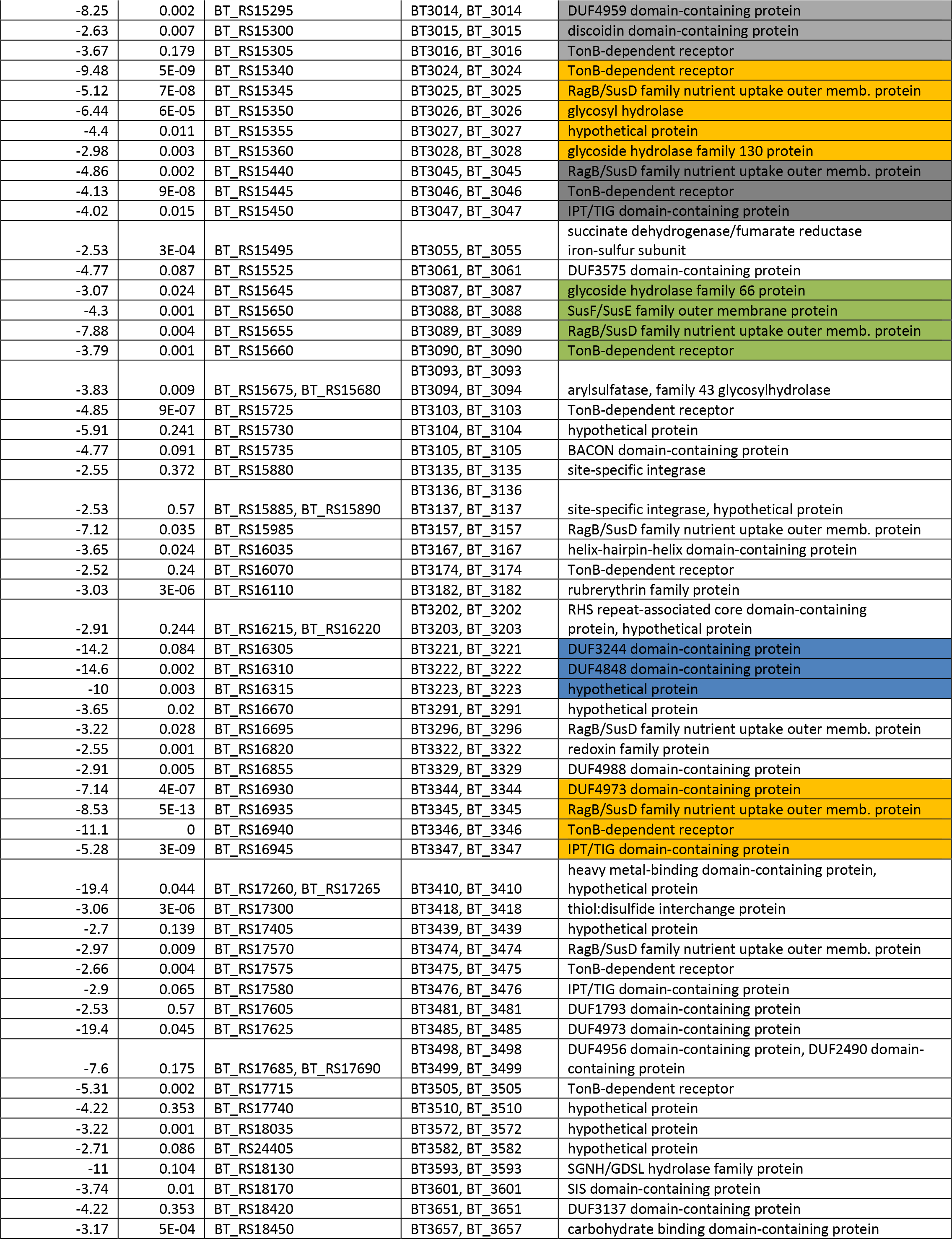
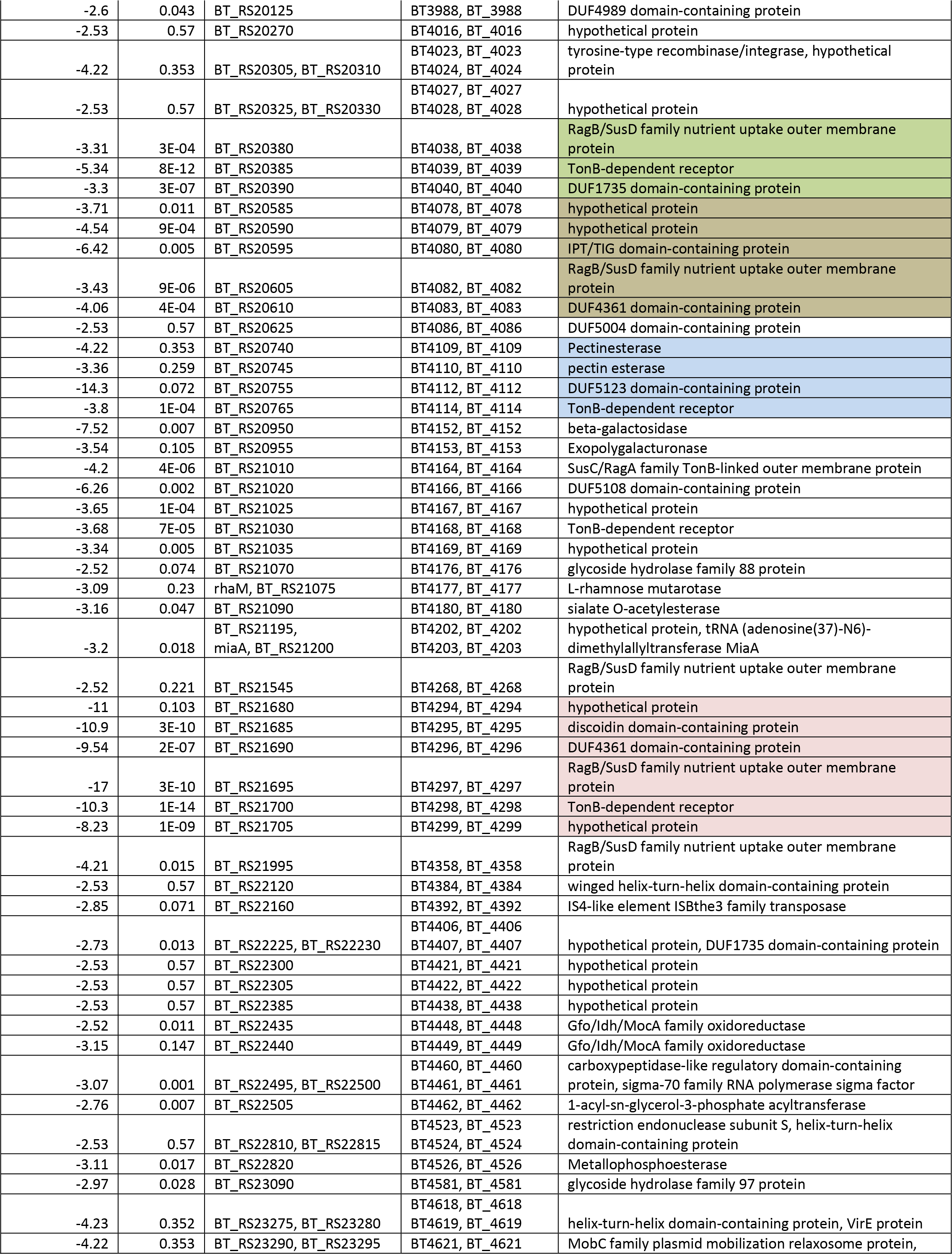
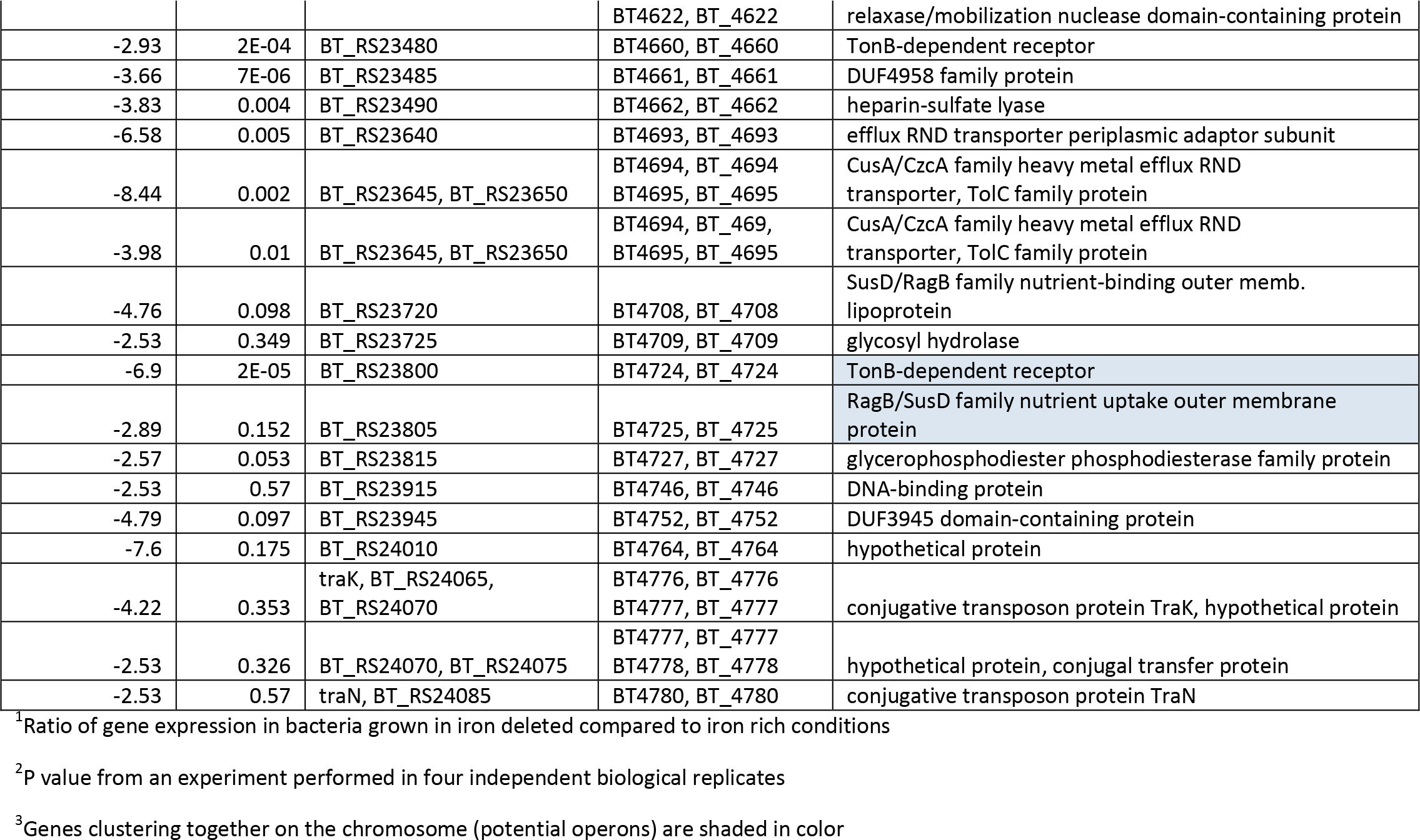
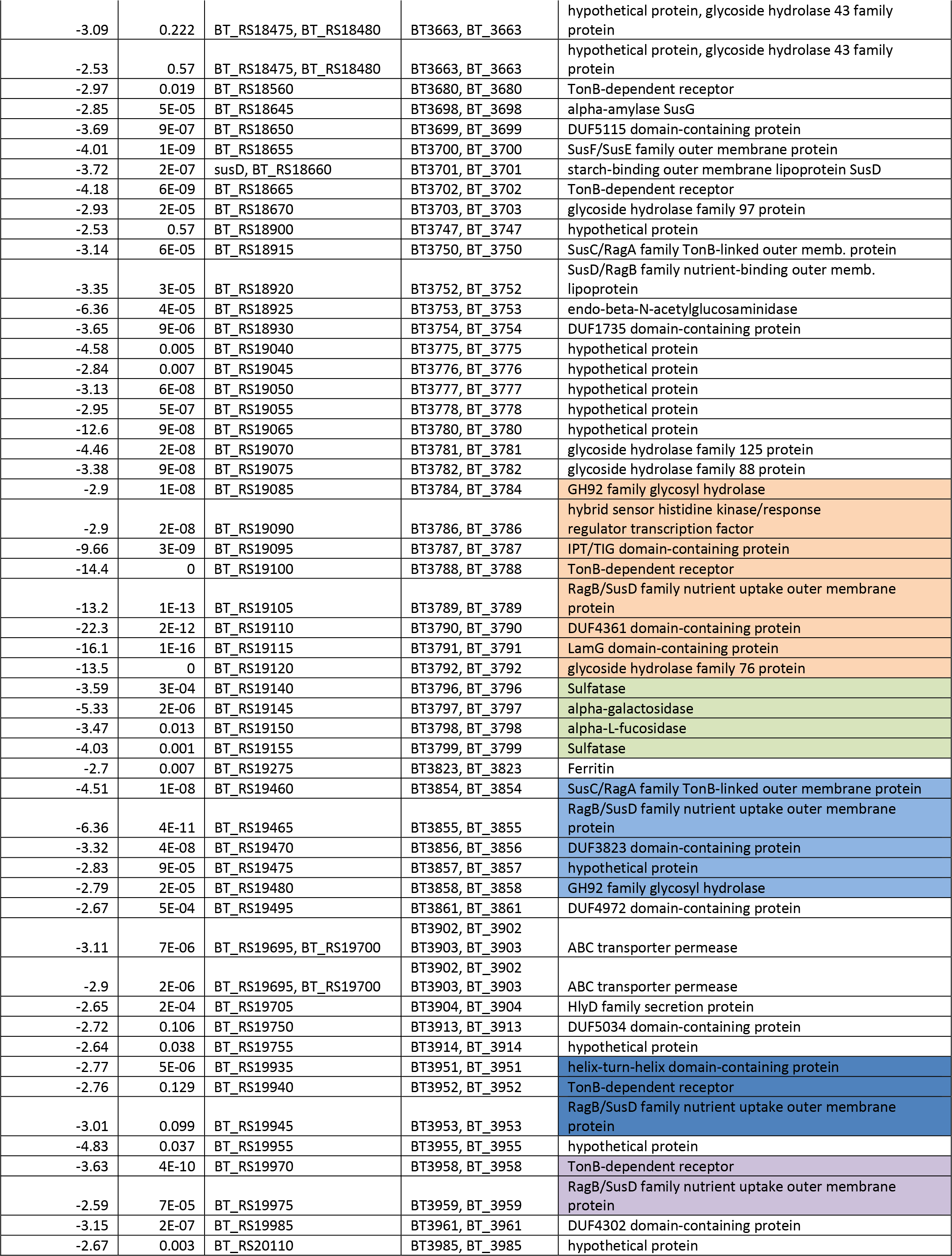
Genes downregulated in B. thetaiotaomicron BT5482 Δ*tdk* at least 2.5 fold under iron deplete conditions.

Two genes, BT0519 and BT0520, coding for hemerythrin domain-containing protein, and a response regulator/transcription factor were upregulated by 16.3 and 20fld, respectively. An operon coding for carbohydrate metabolism composed of BT1658 – BT1660 60 was upregulated by 15.4, 16.8 and 20 fold, respectively. Also, BT2818 – BT2825 coding for a locus containing the SusD/SusC system was upregulated 4.5 –22.5 fold, depending on a gene.

Oxidative stress protection mechanisms such as gene BT0028 coding for alkaline phosphatase was upregulated 15.4 fold. BT2811 coding for alkyl hydroperoxide reductase subunit F was also upregulated by 8 fold. Finally, gene BT4815 coding for the DNA starvation/stationary phase protection protein (Dps) was upregulated by 5.8 fld. Finally, iron-independent metabolism operon BT0516 – 517 coding for a flavodoxin (FldA, BT0517) and DUF2023 protein (BT0516) were upregulated 326 and 598 fold, respectively. This operon was also reported to be highly upregulated in the colitis model^2^ (Table 7).

**Table 7.** Clinical correlation. B. thetaiotaomicron genes regulated by iron and in colitis.

| **-Fe/**<br>**+Fe<sup>1</sup>** | **Gene**<br>**Name<sup>2</sup>** | **Colitis**<br>**Study<sup>3</sup>** | **Protein annotation<sup>4</sup>** | **Presence<sup>5</sup>** |
| --- | --- | --- | --- | --- |
| 1.898914 | BT_0159 | 2.111094 | VIT1/CCC1 transporter family protein | Bt |
| 8.06201 | BT_0224 | 9.704741 | hypothetical protein | Bt |
| 4.40866 | BT_0225 | 11.11973 | hypothetical protein | Bt, Pi |
| 4.883878 | BT_0226 | 10.46389 | hypothetical protein | Bt |
| 3.419034 | BT_0227 | 7.563568 | porin family protein | Bt |
| 1.633861 | BT_0239 | -1.05735 | DUF4369 domain-containing protein | Bt, Pg, Pi |
| 108.9247 | BT_0491 | 18.86303 | DUF2149 domain-containing protein,<br>MotA/TolQ/ExbB proton channel family<br>protein | Bt,, Pg, Pi |
| 186.9863 | BT_0492 | 28.50811 | DUF2149 domain-containing protein,<br>MotA/TolQ/ExbB proton channel family<br>protein | Bt,, Pg, Pi |
| 136.0809 | BT_0493 | 16.425 | hypothetical protein | Bt, , Pg, Pi |
| 282.4144 | BT_0496 | 24.69941 | TonB-dependent receptor | Bt,, Pg, Pi |
| 290.8443 | BT_0497 | 29.40818 | HmuY family protein | Bt |
| 263.7668 | BT_0498 | 31.73038 | hypothetical protein | Bt, Pg, |
| 2.392774 | BT_0503 | 8.93965 | hypothetical protein | Bt,Pg |
| 2.430991 | BT_0504 | 4.850745 | TonB-dependent receptor | Bt, Pg, Pi |
| 51.24636 | BT_0507 | 5.508571 | TetR/AcrR family transcriptional regulator | Bt, Pi, Pi |
| 675.5204 | BT_0508 | 15.40959 | TetR/AcrR family transcriptional regulator | Bt, Pi |
| 516.2514 | BT_0509 | 11.91826 | ABC transporter ATP-binding protein | Bt, |
| 37.04651 | BT_0515 | 3.708254 | quinol oxidase | Bt, Pg, Pi |
| 598.3778 | BT_0516 | 27.22851 | DUF2023 family protein | Bt, Pg, Pi |
| 325.7 | BT_0517 | 29.3 | Flavodoxin | Bt, Pg, Pi |
| 19.95236 | BT_0519 | 9.597707 | hemerythrin domain-containing protein,<br>response regulator | Bt, Pg, Pi |
| 16.32074 | BT_0520 | 9.573912 | hemerythrin domain-containing protein,<br>response regulator transcription factor | Bt |
| 2.014698 | BT_0523 | 4.325417 | sugar transferase | Bt |
| 2.142719 | BT_0524 | 1.586668 | response regulator | Bt, |
| 2.007205 | BT_0532 | 7.851734 | anthranilate synthase component I family<br>protein | Bt, Pg |
| -4.22 | BT_0545 | -174 | P-II family nitrogen regulator | Bt |
| -2.5 | BT0551 | -167.1 | Asparagine synthase | Bt |
| -3.5 | BT0552 | -1267 | Glutamate synthase beta subunit | Bt |
| -1.99 | BT0553 | -290 | Glutamate synthase large subunit | Bt |
| 3.711937 | BT_0922 | 4.113622 | PepSY-like domain-containing protein | Bt, Pg |
| 5.042992 | BT_0923 | 3.241869 | PepSY-like domain-containing protein | Bt |
| 2.027212 | BT_0995 | 1.799071 | PAS domain-containing sensor histidine<br>kinase, helix-turn-helix domain-containing<br>protein | Bt |
| 3.760234 | BT_1043 | 10.62892 | SusD/RagB family nutrient-binding outer<br>membrane protein | Bt |
| 8.318422 | BT_1045 | 10.67078 | DUF1735 domain-containing protein | Bt, Pg, Pi |
| 3.099403 | BT_1138 | -1.56722 | hypothetical protein, site-specific<br>integrase | Bt |
| 2.279712 | BT_1597 | 3.318077 | tetratricopeptide repeat protein, HAMP<br>domain-containing histidine kinase | Bt |
| 3.294873 | BT_1895 | 35.68394 | hypothetical protein | Bt |
| 3.128725 | BT_1896 | 40.56019 | leucine-rich repeat protein | Bt, Pg |
| 3.320197 | BT_2038 | 5.169461 | efflux RND transporter periplasmic<br>adaptor subunit | Bt, Pg |
| 2.477183 | BT_2039 | 6.533055 | CusA/CzcA family heavy metal efflux RND transporter | Bt, Pg |
| 2.549452 | BT_2040 | 4.430047 | TolC family protein | Bt |
| 489.3635 | BT_2063 | 34.05744 | PepSY domain-containing protein | Bt |
| 676.0915 | BT_2064 | 34.84073 | DUF4374 domain-containing protein | Bt, Pi |
| 662.945 | BT_2065 | 16.46505 | TonB-dependent receptor | Bt, Pi |
| 3.463556 | BT_2450 | 26.23424 | hypothetical protein | Bt |
| 62.68564 | BT_2473 | 14.21087 | fimbrillin family protein | Bt |
| 103.9787 | BT_2475 | 29.12251 | cytochrome c biogenesis protein CcsA, hypothetical protein | Bt |
| 225.3287 | BT_2476 | 28.66125 | hypothetical protein | Bt |
| 146.1537 | BT_2477 | 25.62196 | porin | Bt |
| 133.2681 | BT_2478 | 30.08073 | thiol oxidoreductase | Bt |
| 230.1729 | BT_2479 | 32.79333 | peptidase M75 | Bt |
| 283.9951 | BT_2480 | 26.77281 | hypothetical protein | Bt |
| 323.7068 | BT_2482 | 20.39152 | helix-turn-helix transcriptional regulator | Bt |
| 2.264999 | BT_2778 | 1.409648 | sigma-70 family RNA polymerase sigma factor | Bt |
| -2.26543 | BT_3053 | -3.48253 | succinate dehydrogenase/fumarate reductase cytochrome b subunit | Bt, Pg, Pi |
| -2.29584 | BT_3054 | -2.86087 | fumarate reductase/succinate dehydrogenase flavoprotein subunit | Bt, Pg, Pi(p) |
| 65.02367 | BT_3625 | 15.21476 | DUF4857 domain-containing protein | Bt |
| 36.69843 | BT_3626 | 14.07917 | hypothetical protein | Bt |
| 39.85516 | BT_3627 | 18.25618 | ABC transporter ATP-binding protein | Bt, |
| 38.27088 | BT_3628 | 17.59845 | PepSY-associated TM helix domain-containing protein | Bt |
| 43.33606 | BT_3629 | 17.27476 | PepSY-associated TM helix domain-containing protein | Bt |
| 97.71608 | BT_3630 | 19.14698 | fimbrillin family protein | Bt |
| 128.0625 | BT_3631 | 14.74463 | hypothetical protein, DUF4876 domain-containing protein | Bt |
| 125.3197 | BT_3632 | 16.11913 | hypothetical protein, DUF4876 domain-containing protein | Bt |
| 112.1206 | BT_3633 | 18.93561 | TonB-dependent receptor | Bt |
| 5.035874 | BT_3969 | 3.185176 | CusA/CzcA family heavy metal efflux RND transporter | Bt |
| 2.017194 | BT_4227 | 13.80192 | Mfa1 fimbrillin C-terminal domain-containing protein | Bt, |
| 2.305598 | BT_4233 | 9.275554 | DUF3845 domain-containing protein | Bt |
| 3.306148 | BT_4234 | 9.429591 | DUF3575 domain-containing protein | Bt, Pi |
| 2.182647 | BT_4236 | -1.25097 | response regulator | Bt |
| 4.047878 | BT_4571 | 3.342238 | hypothetical protein, RNA polymerase sigma factor | Bt |
| 5.844437 | BT_4715 | 18.7437 | DNA starvation/stationary phase protection protein | Bt, Pg, Pi |

<sup>1</sup> – fold change in this study (-Fe/+Fe)
<sup>2</sup> – name of *B. thetaiotaomicron* VPI – genes
<sup>3</sup> – fold change in colitis study<sup>2</sup>
<sup>4</sup> – protein annotation
<sup>5</sup> – presence of the gene in *B. thetaiotaomicron* (Bt), *P. gingivalis* (Pg), *P. intermedia* (Pi)

Downregulated 2.5 fold were 444 genes. The regulated genes were arranged into 51 operons. Among the most drastically downregulated operons was the operon coding for dissimilative nitrite reduction to ammonia BT1414 – 418 that includes the *nrfAH* (BT1417-18), cytochrome reductase preceded by the cytochrome c biogenesis coding genes *ccsA* (BT1415) and *ccsB* (BT1416)(Table 6, Fig. 3I). That operon also included a gene coding for an alginate export family protein (BT1414). It is predicted to be regulated by a Crp-like regulator encoded by BT1413.

To extend the nitrogen regulation cycle, downregulated was also gene coding for nitrogen-fixing protein NifU (BT0174) and *hcp* (BT0687) coding for putative hydroxylamine reductase, Hcp (downregulated [-3.6 fold]). The gene adjacent to *hcp* coding for uracil-xanthine permease (BT0686) was also downregulated by 2.5 fold suggesting the two genes form an operon. *hcp* is predicted to be regulated by HcpR which is encoded by BT0688.

Genes encoding iron-dependent metabolic mechanisms including the succinate dehydrogenase/fumarate reductase (BT3055) and RnfABCDGE electron transport complex (BT0618 – 622) was downregulated by 2.5-3.4 fold, depending on a gene. Gene encoding asparagine biosynthesis, BT0551 was downregulated by 2.5 and a locus coding for glutamate synthase (BT0552 -3) was downregulated by 3.5 fold (Fig. 3L). Finally, locus *hydGEF* (BT1834-837) coding for a 4Fe-4S cluster containing hydrogenase was downregulated by 2.9 – 3.6 fold, depending on a gene.

Iron homeostasis and oxidative stress encoding genes including ferritin coding gene (*ftn,* BT3823) and rubrerythrin family protein coding gene (BT3182) were downregulated by 2.7 – 3 fold, depending on a gene (Table 6). Of interest was also locus encoding sulfatases, BT3796 – BT3799, downregulated by 3.5-5.3 fold depending on a gene (Table 6).

Large number of operons coding for two-component Sus-like transporters systems that contained the gene coding for TonB-dependent receptor as well as the Rag/Sus family nutrient uptake protein (BT4724-25, BT4294-99, BT4038-40, BT3958-59, BT3951-53, BT3854-58, BT3786-92, BT3344-47), BT3087-90, BT3045-47, BT3024-28, BT3012-16, BT2804-09, BT2614-32, BT2392-95, BT1280-85, BT1046-47, BT0438-40, BT0360-65, BT0317-18) were downregulated, Table 6. Finally, conjugative transposon coding genes *traN, traM,* and *traK* (BT0086-89) were also downregulated by 2.5-7.6 fold, depending on a gene.

Overall, iron/hemin uptake, carbohydrate metabolism, oxidative stress and iron-independent metabolism were upregulated. Downregulated were genes that code for proteins containing iron cofactors such as metabolic enzymes (*nrfAH, ccsA, hcp, rnf, hyd*), oxidative stress (*rbr, ftn*), as well as TonB and RagA/Sus transport systems. Lower iron levels also reduced expression of genes involved in DNA mobilization.

### Iron-dependent stimulon of Bacteroidetes

Comparison of the regulated genes across the three bacterial species analyzed in our study has identified many common genes (Fig. 4) (similarity in gene sequence and predicted functions) (Figs. 1 – 3). Furthermore, many of the genes had similar genomic organization in the three bacterial species studied. Among the commonly regulated operons was the *hmu* encoding hemin update mechanism (Tables 1, 3, and 5, Fig. 1A). Also, the flavodoxin-encoding gene was highly upregulated in all three bacterial species tested (Fig. 1B). The locus is composed of two genes: *fldA* and a downstream gene coding for a hypothetical protein. In *B. thataiotaomicron* the *fldA* and the downstream gene, BT0517 and BT0516, are predicted to form an operon. Such prediction is further substantiated by the significant upregulation of both genes (Table 5). Since both genes are also present in the oral bacteria it is likely that those are also co-transcribed. Two operons containing TonB – dependent receptors were upregulated in three (Fig. 1C), and two (Fig. 1D) bacteria, respectively. Drastically upregulated in oral species, *P. gingivalis* and *P. intermedia* was PG0495 (upregulated 87fld) and PIOMA14_RS06105 (ChI, upregulated 104 fld) (in *P. intermedia* 17: PIN17_RS05350) (Fig. 1E, Supplemental Fig. 1). This gene encodes a T9SS C-terminal target domain protein. Of note, in *P. intermedia* this locus is located next to *hmuY* (Fig. 1E).

**Figure 1.**
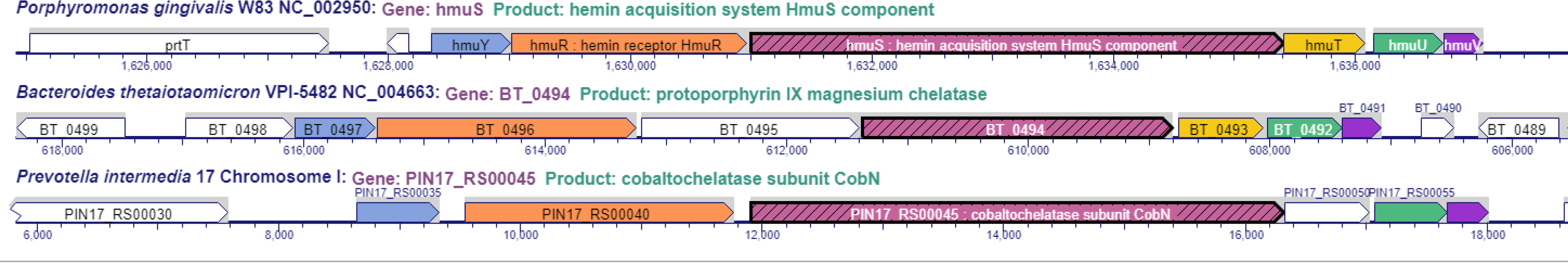
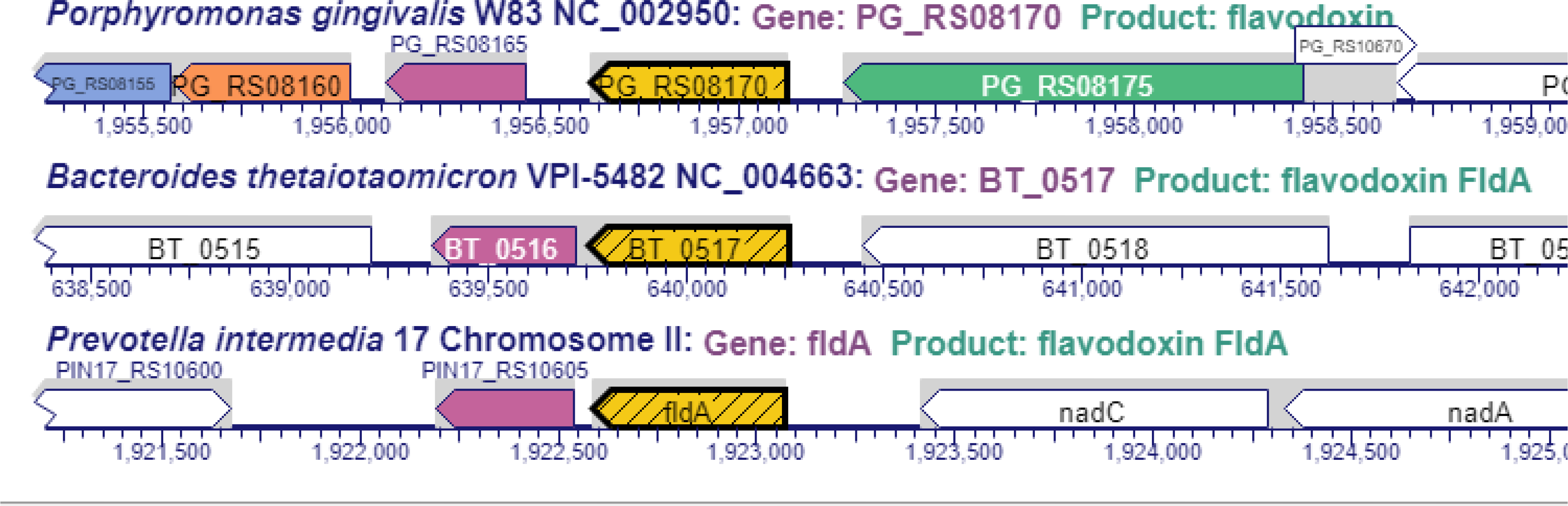
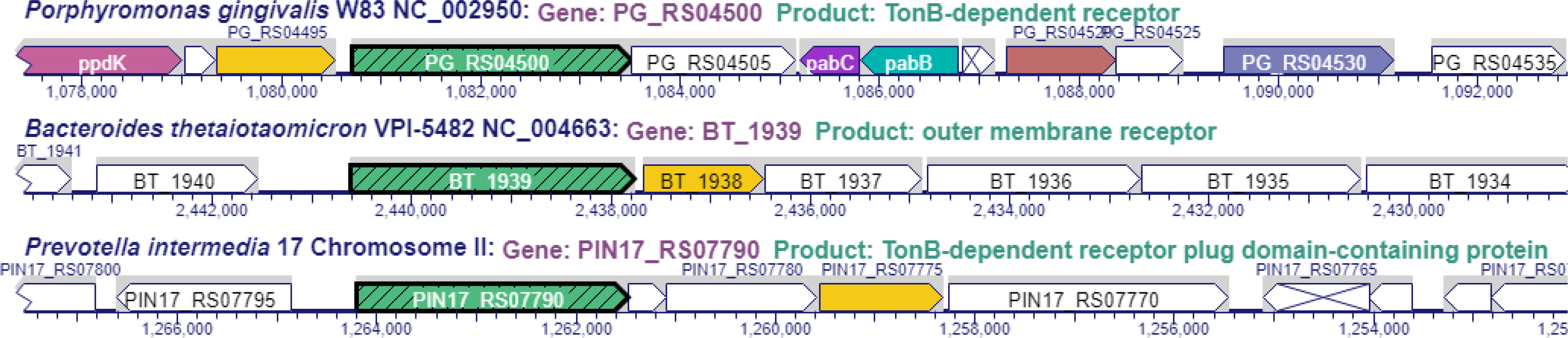
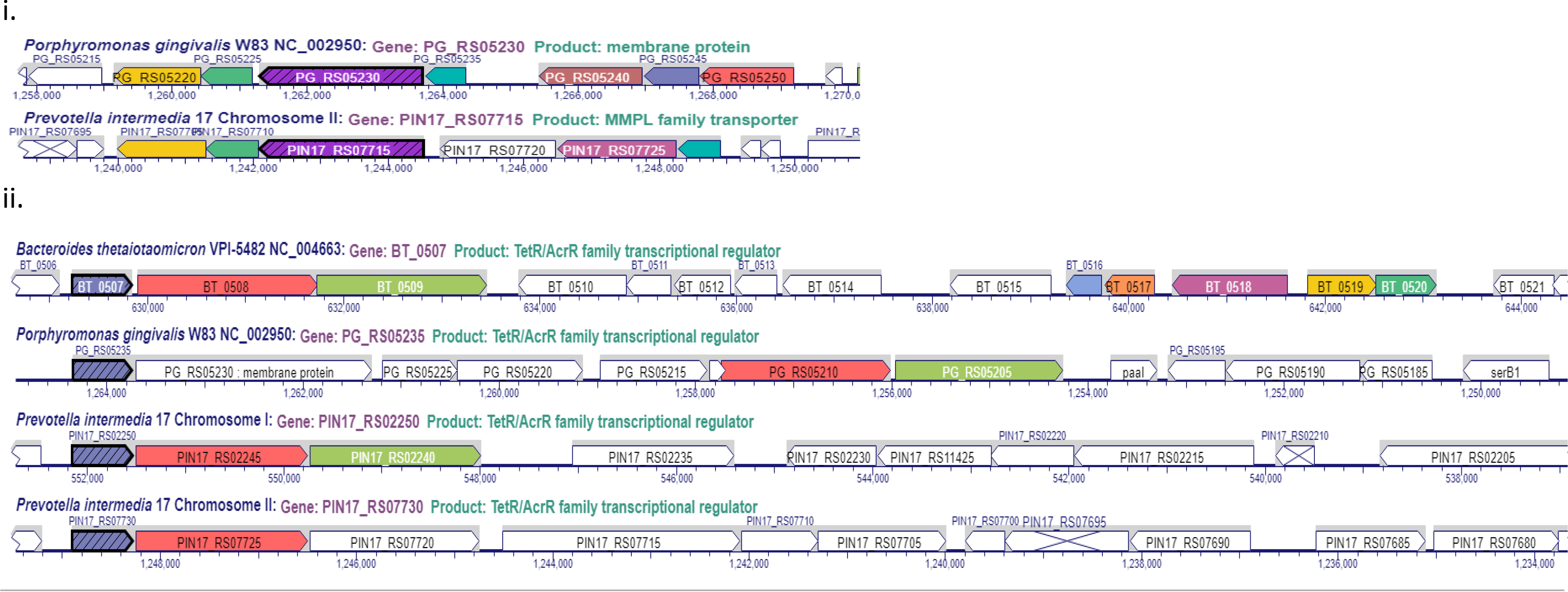
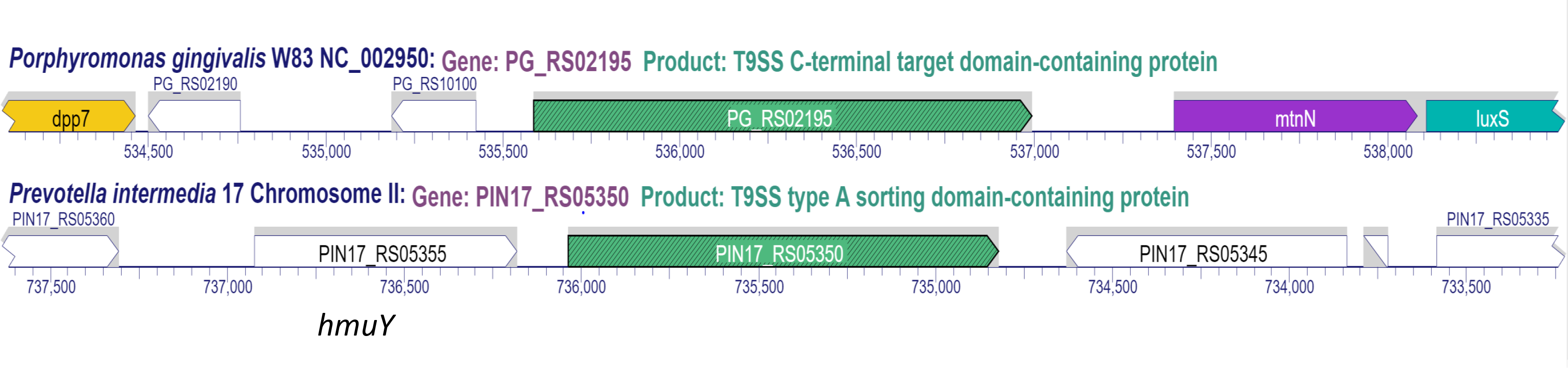
Comparison of loci upregulated in Bacteroides. A. Hemin uptake *hmu* locus in Bacteroidetes: comparison of genomic locus in *P. gingivalis* W83 (*hmuY* – *hmuT*, PG_RS0), *B. thetaiotaomicron* VPI-5482 (BT_0491 -98), *P. intermedia* 17 (PIN17_RS00035-55). B. Flavodoxin *fld* locus in Bacteroidetes (designated in yellow): comparison of genomic locus *P. gingivalis* W83 (PG_RS08170, PG1858), *B. thetaiotaomicron* VPI-5482 (BT_0517), *P. intermedia* 17 (*fldA*). C. Outer membrane transport system locus in Bacteroidetes: comparison of genomic locus *P. gingivalis* W83 (PG_RS04495 – PG_RS04505; PG1019-21), *B. thetaiotaomicron* VPI-5482 (BT_1938-9), *P. intermedia* 17 (PIN17_RS07775 – 90). D. TetR/AcrR regulator and transport system locus in Bacteroidetes: comparison of genomic locus *P. gingivalis* W83 (PG_RS05220 – 235; PG1178 - 81), *B. thetaiotaomicron* VPI-5482 (BT_0507-9), *P. intermedia* 17 (PIN17_RS07705 – 25, PIN17_RS02240 50, PIN17_RS07725-30).E. Cell surface – extracellular protein in Bacteroidetes: comparison of genomic locus *P. gingivalis* W83 (PG_RS02195, PG0495), *B. thetaiotaomicron* VPI-5482 (no ortholog found), *P. intermedia* 17 locus (PIN_RS05350).

There were also several commonly downregulated genes. The iron-based metabolism coding genes including the operon *sdhABC* coding for succinate dehydrogenase/fumarate reductase (Fig. 2A), the *rnf* coding for the electron transport chain (Fig. 2B), *vor,* the 2-oxoglutarate ferredoxin oxidoreductase encoding genes (Fig. 2C) were downreglated in all three bacterial species tested. Rubrerythrin-encoding gene, *rbr,* was more drastically downregulated in *P. gingivalis* and *P. intermedia* (PG0195 and PIOMA14_II_0105 with 46 and 22.3fold change, respectively) compared to *B. thetaiotaomicron* (BT3812 downregulated by 3 fold) (Fig. 2D, Tables 2, 4, and 6). It is noteworthy that *B. thetaiotaomicron* encodes another rubrerythrin (BT0216) that is proceeded by BT0215 coding for a putative iron uptake regulatory protein, Fur, but that locus was not significantly affected by iron. Also, the *ahp* locus coding for the alkylhydroperoxide reductase (Fig. 3B) was found to be upregulated in *P. gingivalis* and *B. thetaiotaomicron*.

**Figure 2.**
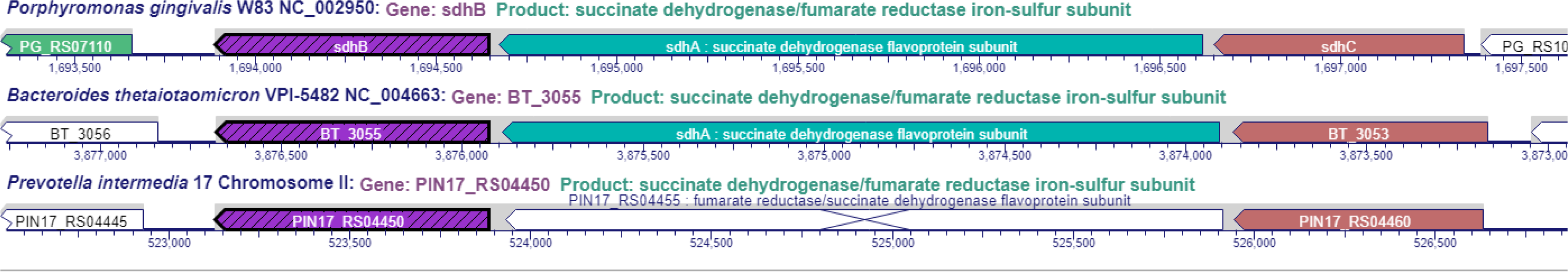
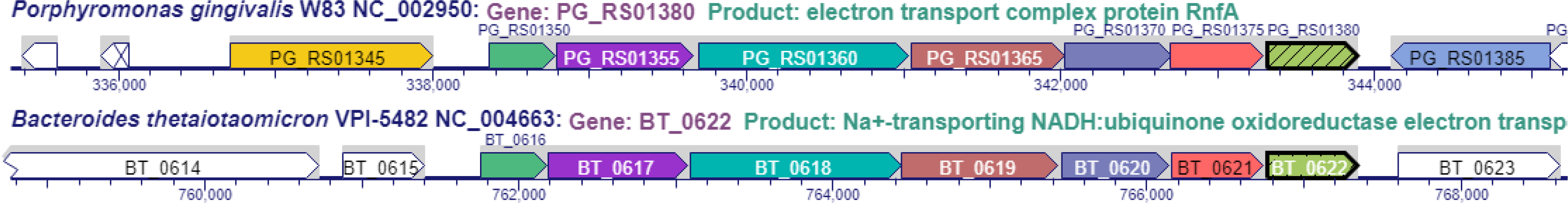
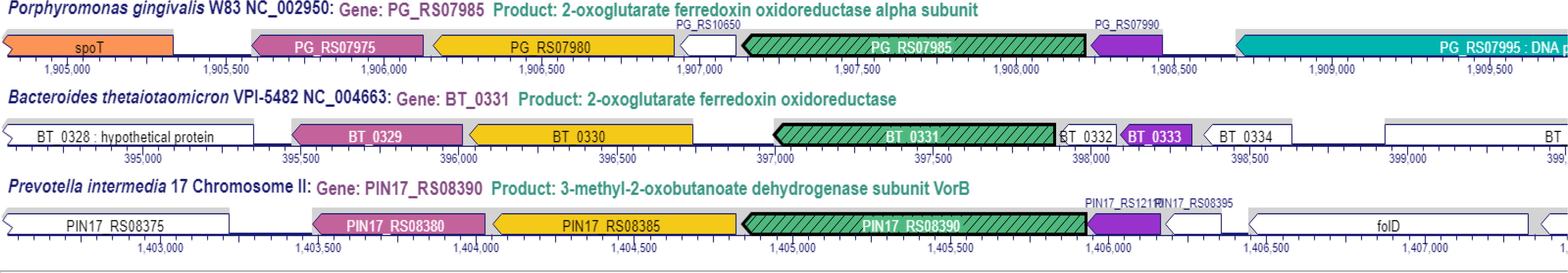
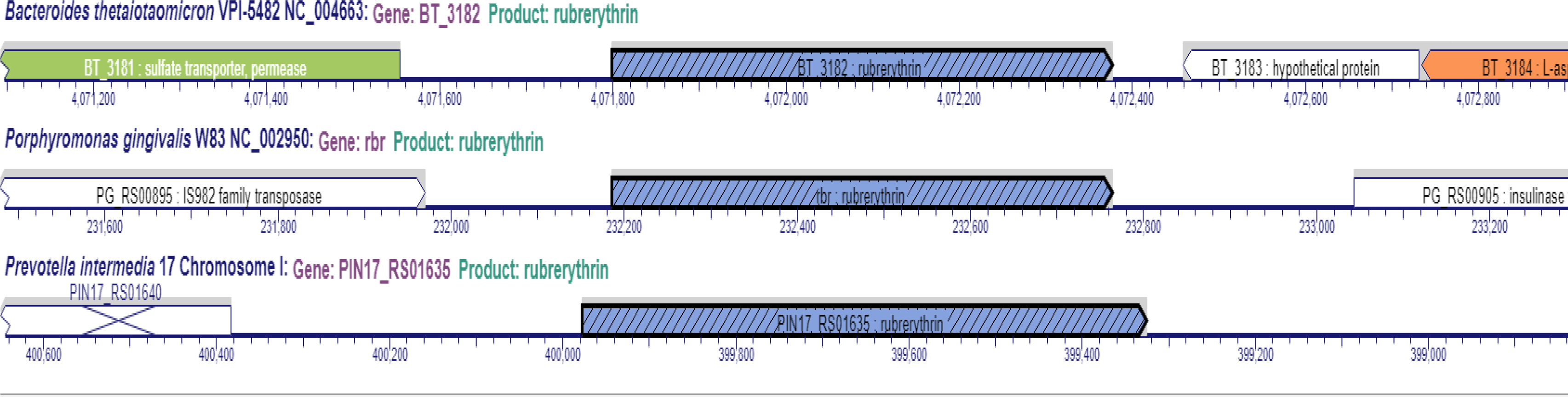
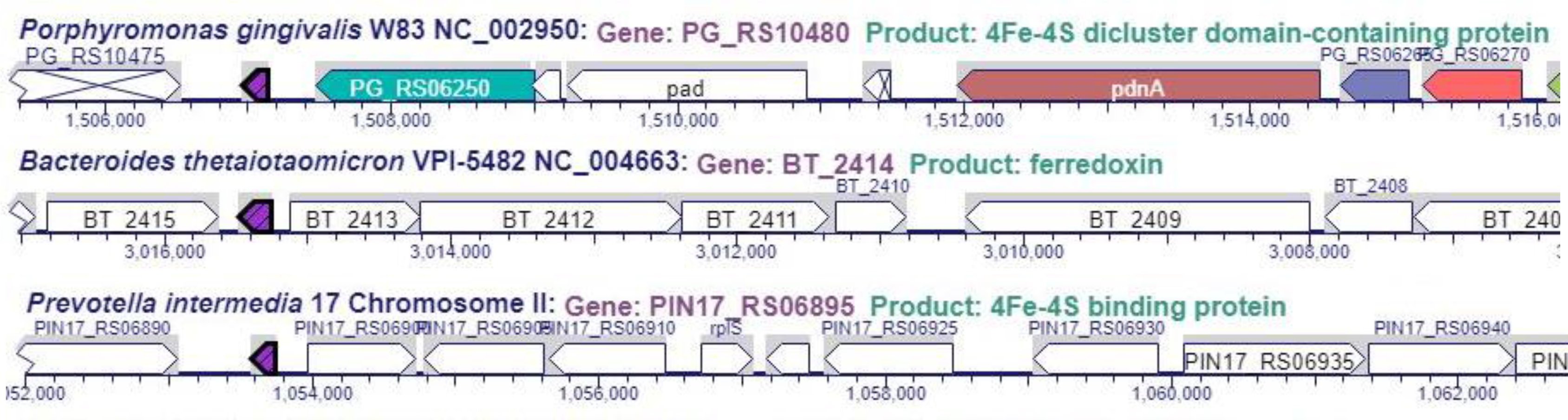
Comparison of loci downregulated in Bacteroides. A. Succinate dehydrogenase/fumarate reductase *sdh* locus locus in Bacteroidetes: comparison of genomic locus of *P. gingivalis* W83 (PG_RS07115 – 125, PG1614-16), *B. thetaiotaomicron* VPI-5482 (BT_3053-55), *P. intermedia* 17 (PIN17_RS04460 – 50). B. Electron transport *rnf* locus locus in Bacteroidetes: comparison of genomic locus of *P. gingivalis* W83 (PG_RS01350 -80, PG0302 – 308), *B. thetaiotaomicron* VPI-5482 (BT_0617 – 22). No ortholog of the locus was not found in *P. intermedia* 17. C. Ferredoxin- oxidoreductase (*vor*) locus locus in Bacteroidetes: comparison of genomic locus of *P. gingivalis* W83 (PG_RS07975-90, PG1809013), *B. thetaiotaomicron* VPI-5482 (BT_0329 – BT_0333), *P. intermedia* 17 (PIN17_RS08380 – 95). D. Rubrerythrin (*rbr*) locus locus in Bacteroidetes: comparison of genomic locus of *P. gingivalis* W83 (PG_RS00900, PG0195), *B. thetaiotaomicron* VPI-5482 (BT_3182), *P. intermedia* 17 (PIN17_RS01635). E. 4Fe-4S dicluster domain-containing protein (frd) locus locus in Bacteroidetes (purple arrow): comparison of genomic locus of *P. gingivalis* W83 (PG_RS10480, PG1421), *B. thetaiotaomicron* VPI-5482 (BT_2414), *P. intermedia* 17 (PIN17_RS06895).

**Figure 3.**
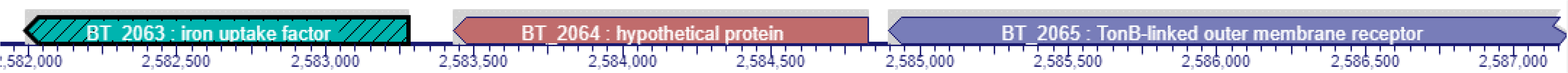
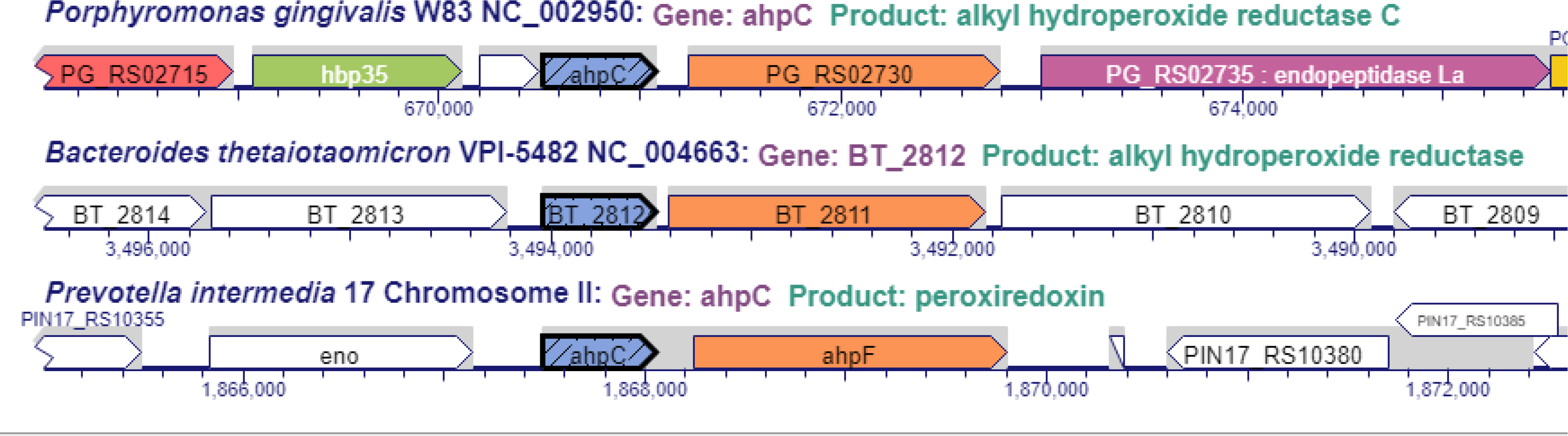
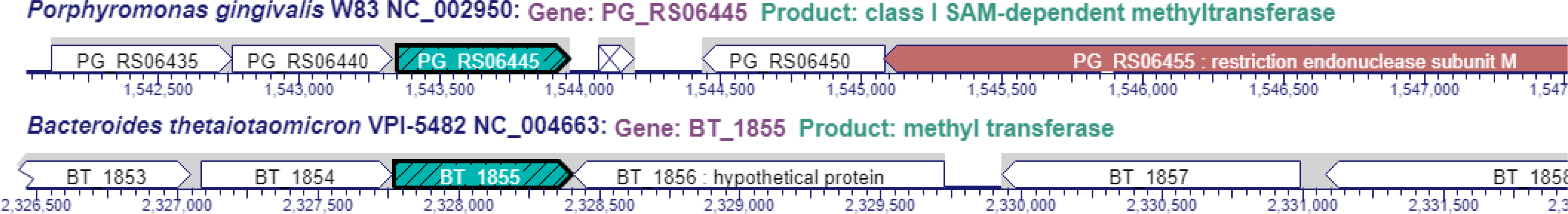
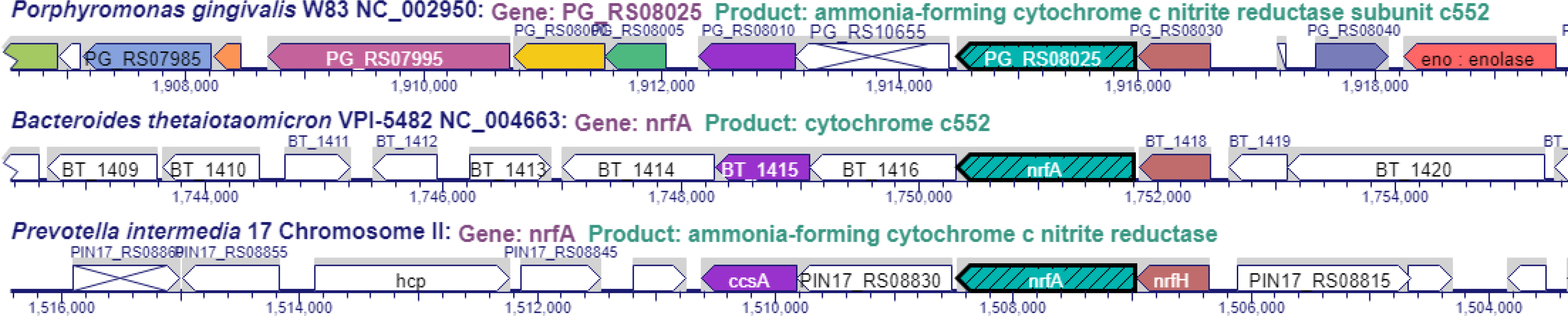
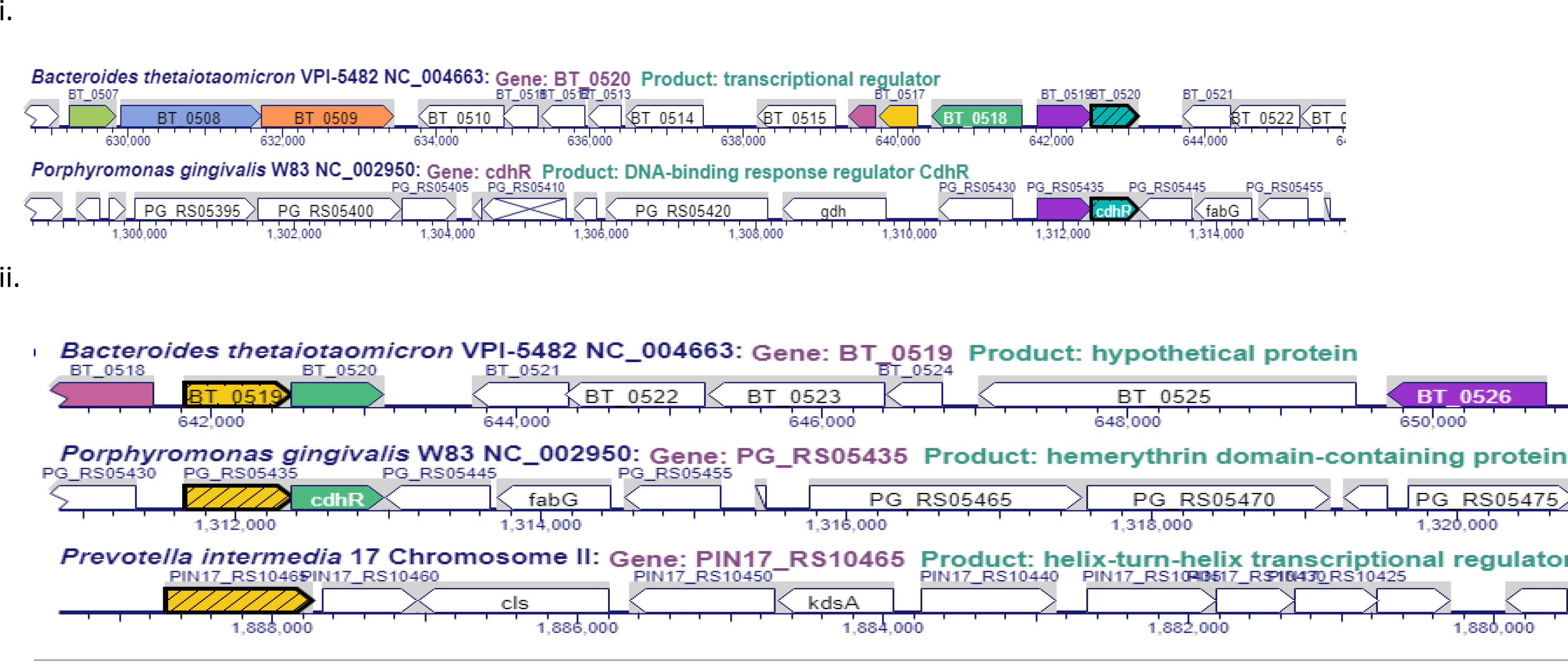
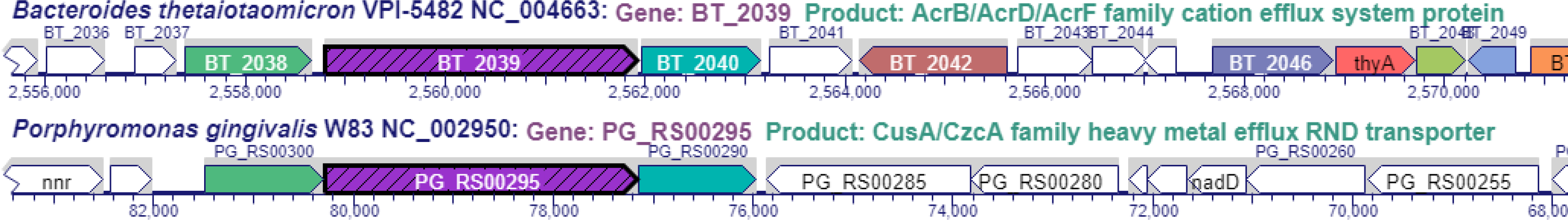
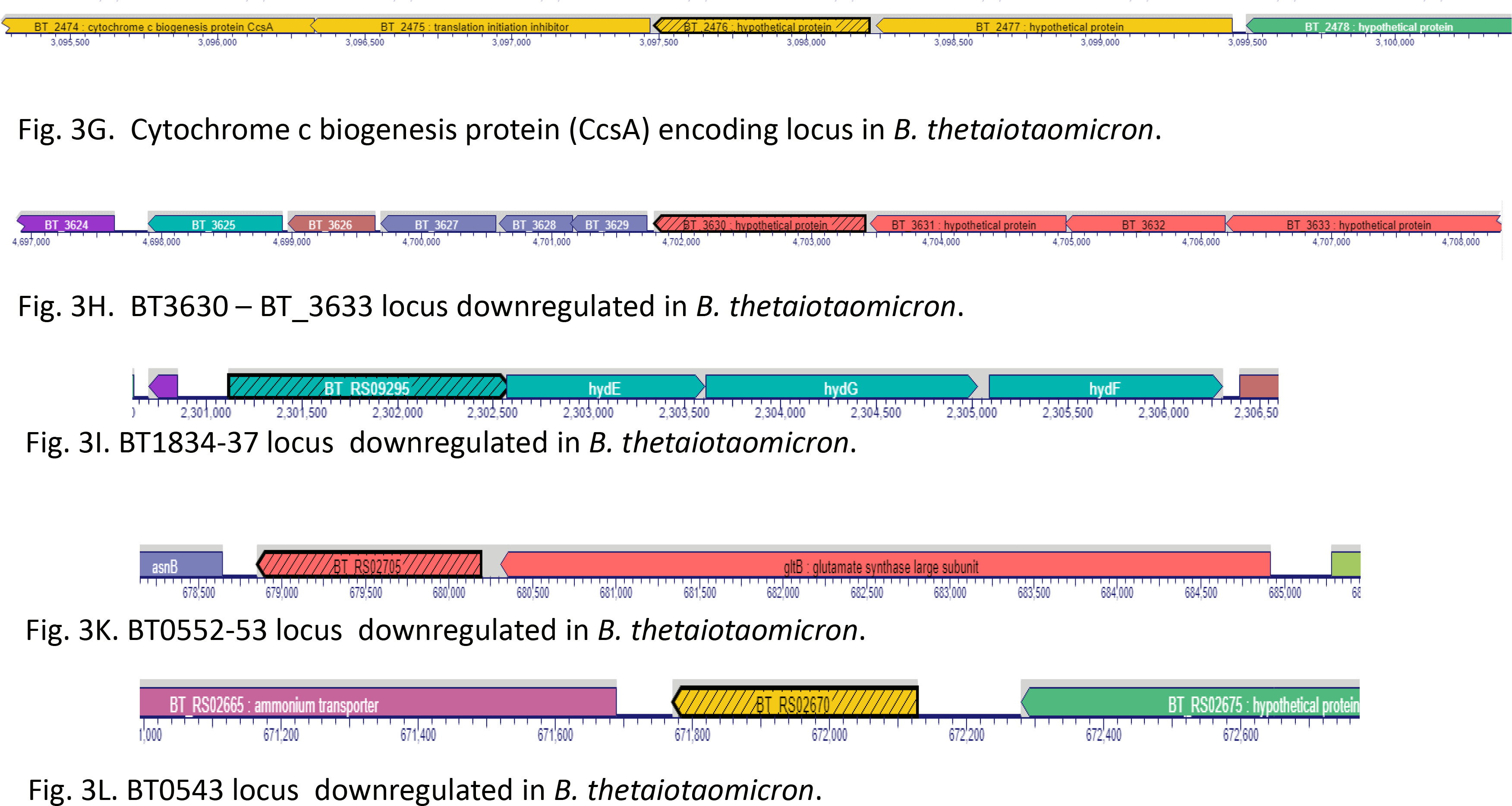
Comparison of other loci regulated by iron in Bacteroidetes. A. Xenosiderophore-mediated iron acquisition encoding locus in *B. thetaiotaomicron* (BT_2063 – 2065). The BT_2065 coding for the TonB-outer membrane receptor has an ortholog in *P. intermedia.* B. *ahpCF* locus locus in Bacteroidetes: comparison of genomic locus of *P. gingivalis* W83 (new locus tag: PG_RS02725 – 30. Old locus tag: PG0618 – 19), *B. thetaiotaomicron* VPI-5482 (BT2811-12), *P. intermedia* 17 (*ahpC-ahpF*). C. SAM-methyltransferase encoding locus in Bacteroidetes: comparison of genomic locus of *P. gingivalis* W83 (new locus tag: PG_RS06445. Old locus tag: PG1467), *B. thetaiotaomicron* VPI-5482 (BT_1855). No similar locus was identified in *P. intermedia* 17. D. *nrfAH* locus in Bacteroidetes: comparison of genomic locus of *P. gingivalis* W83 (new locus tag: PG_RS0825 – 30. Old locus tag: PG1820-21), *B. thetaiotaomicron* VPI-5482 (BT_1417-18), *P. intermedia* 17. E. Hemerythrin-domain containing regulator (CdhR in *P. gingivalis*). i. identical operon coding for an hemerythrin-domain followed by a DNA- binding regulator (CdhR) is present in *P. gingivalis* (new locus tag: PG_RS05435 – cdhR. Old locus tag: PG1236-37) and *B. thetaiotaomicron* (BT_0519 – 520). ii. In *P. intermedia* the operon is fused thus producing one regulatory PIN17_RS10465 protein. F. Cation efflux system present in *B. thetaiotaomicron* (BT_2038 – 2040) and *P. gingivalis* (new locus tag: PG_RS00295, PG_RS00300, and PG_RS00290. Old locus tag: PG0063-65). G. BT_2063 – 2065. Xenosiderophore-mediated iron acquisition encoding locus in *B. thetaiotaomicron*. H. Cytochrome c biogenesis protein (CcsA) encoding locus in *B. thetaiotaomicron*. I. BT_3630 – BT_3633 locus in *B. thetaiotaomicron*. J. BT1834-37 locus downregulated in *B. thetaiotaomicron*. K.BT0552-53 locus downregulated in *B. thetaiotaomicron*. M. BT0543 locus downregulated in *B. thetaiotaomicron*.

Among other significantly regulated genes was the BT2063-5 locus encoding the xenosiderophore acquisition mechanism ^2^. That locus is only present in *B. thataiotaomicron* and is not found in *P. gingivalis* and *P. intermedia* 17 (Fig. 3A). Also, present only in *B. thetaiotaomicon* was the locus BT0552-3 coding for glutamate synthase as well as the BT1834-7 encoding hydrogenase system. Another locus of interest, methyltransferase encoding locus was upregulated (Fig. 3C) but only in *P. gingivalis* W83. Finally, *nrfAH* locus encoding nitrite reductase was found in all three bacteria tested (Fig. 3D). However, it was found to be drastically affected by iron depletion only in *B. thetaiotaomicron* (Table 6). Of interest was the family of DUF1661 domain protein encoding genes that were present only in *P. gingivalis* (Supplemental Fig. 4). This is a family of genes coding for small proteins that may have unique functions in adaptation of *P. gingivalis* to iron depletion.

**Figure 4.**
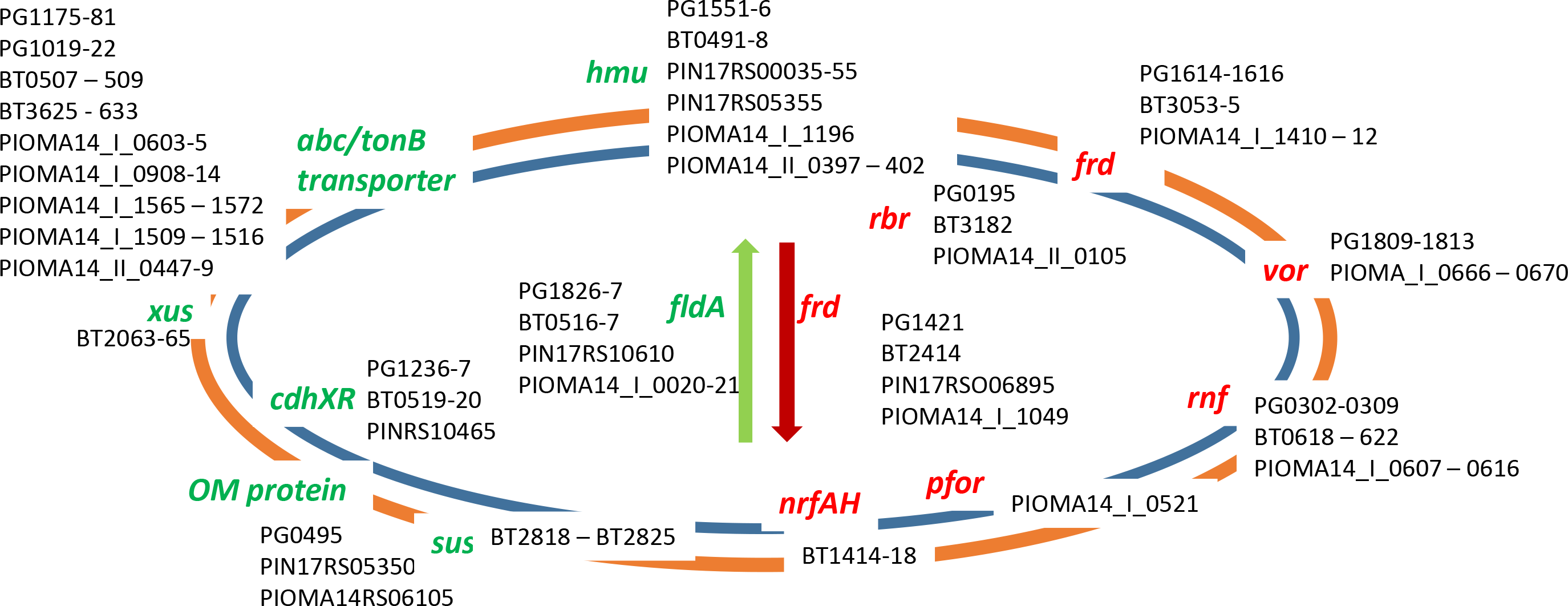
Schematic representation of iron-dependent gene regulation in Bacteroidetes. In bacteria grown under iron limited conditions genes that are upregulated are shown in green while the downregulated ones are shown in red.

Overall, very striking was the drastic upregulation of flavodoxin-encoding gene with concomitant reduction of ferredoxin encoding genes. Consistently with, many iron-based metabolic proteins were downregulated. Upregulated were iron uptake mechanisms possibly to alleviate the iron deficit in the cell (Fig. 4) *Nitrite reduction in B. thetaiotaomicron decreases under iron-limiting conditions.* While all the investigated bacterial species carry genes coding for nitrite reduction, the ammonia-forming cytochrome c nitrite reductase system NrfAH (Fig. 3D), the ability of the bacteria to utilize nitrite has not been demonstrated. Here we show that all three bacterial species reduce nitrite levels present in the culture mixture (Fig. 5AB). This ability differs among the species where *B. thetaiotaomicron* is the most efficient in nitrite reduction. It is able to reduce 2mM nitrite present in culture medium to undetectable levels within 48hr (Fig. 5B). *P. intermedia* OMA14 ranked 2^nd^ in its ability to reduce nitrite while *P. gingivalis* W83 was observed to complete reduction of 0.3mM nitrite to undetectable levels within 72hr (Fig. 5A). While iron depletion decreased the ability of *B. thetaiotaomicron* to reduce nitrite (P=0.065 at 48hr time point) (Fig. 5B) it had no effect on nitrite reduction by *P. gingivalis* and *P. intermedia* (results not shown). The later further functionally verifies the data obtained through our transcriptome sequencing analysis where significant reduction in *B. thetaiotaomicron* transcript levels is observed under iron limiting conditions.

**Figure 5.**
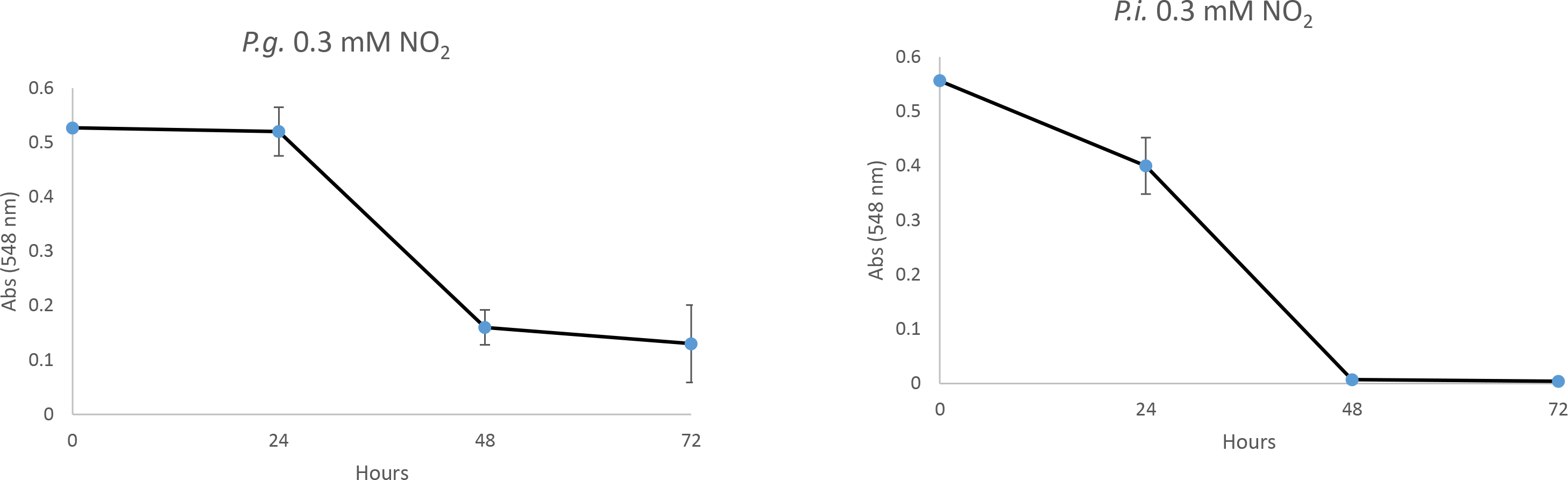
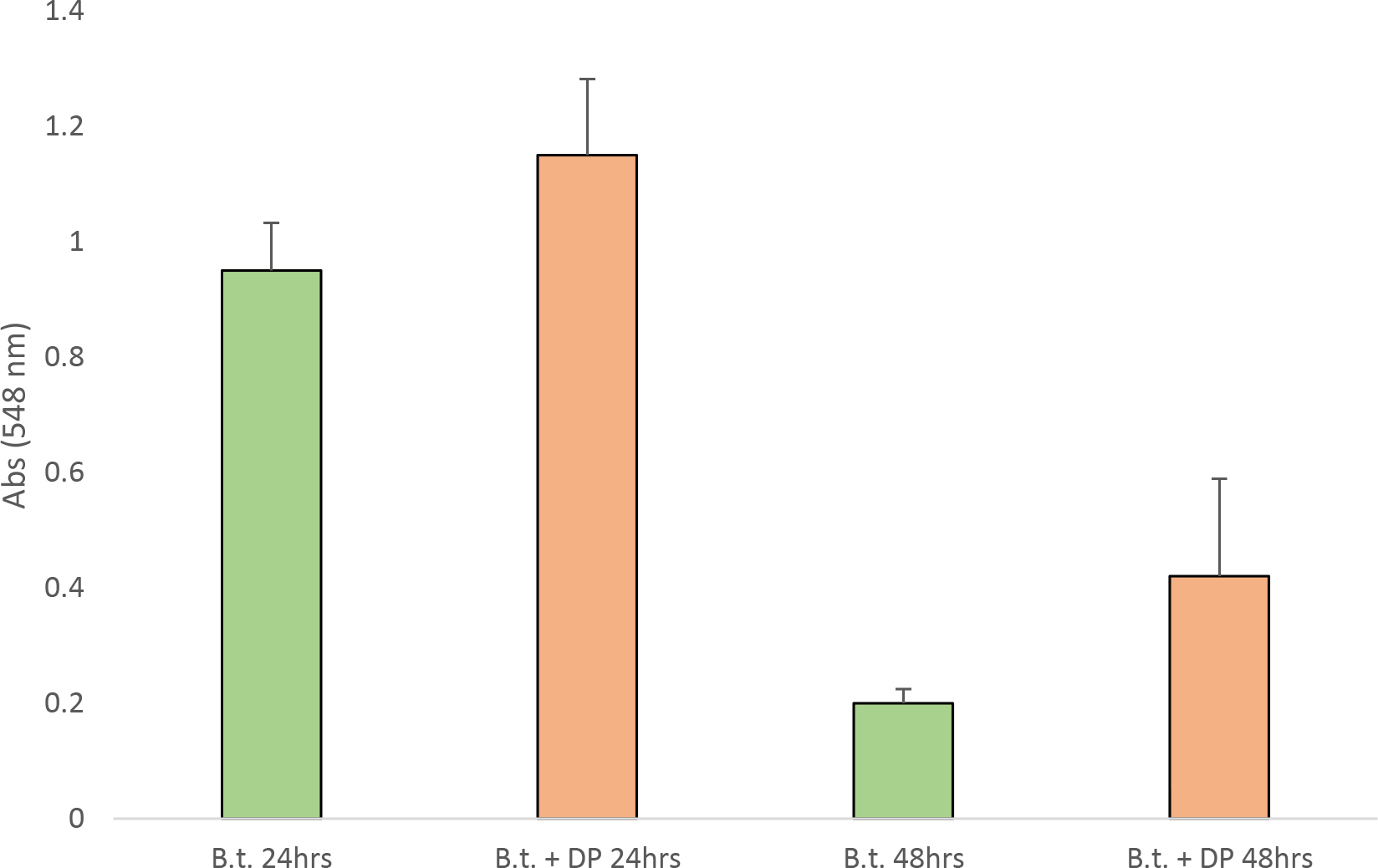
Bacteroides have the ability to reduce nitrite. A. *B. thetaiotaomicron VPI* VPI BT5482 Δ*tdk* was grown in the presence and absence of 2 mM sodium nitrite in TSB medium with and without 150µM of DP. B. *P. gingivalis* W83 and *P. intermedia* OMA14 were grown in the presence and absence of 0.3mM of sodium nitrite in BHI medium. Growth was monitored by measuring optical density at OD_660_ and levels of nitrite were determined with the Griess reagent.

### Clinical correlation of iron-dependent regulation in Bacteroidetes

There was a significant overlap between genes upregulated in *B. thetaiotaomicron* during colitis^2^ and genes upregulated in iron deficient conditions in our study (Table 7). Sixty-eight genes where many were organized into operon structures (15 operon-like organizations) were identified to overlap between the two studies (Table 7). Several loci were present in all three bacteria tested.

First, the locus coding for a flavodoxin (FldA – BT0517) and DUF2023 protein (BT0516) was drastically upregulated in both, our model and the colitis model (Tables 5, 7) ^2^. The *P. gingivalis* flavodoxin encoding gene, *fldA,* was found to be one of the most highly expressed genes in microbiomes derived from human subjects with periodontitis indicating that in the extracellular inflamed environment genes upregulated by iron deficiency are also highly expressed ^24^.

Also significantly upregulated locus both, in our study and during colitis is the *hmu* locus (Table 7). Furthermore, the BT0507 – BT0509 locus coding for TetR/AcrR family transcriptional regulator and two ABC transporter ATP-binding proteins were also upregulated in our study as well as during colitis (Table 7, Fig. 1D). It is noteworthy that two copies of similar operon were found on the genomes of *P. intermedia* (while the copy on the small chromosome was identical to that of the other bacteria studied), the copy on the large chromosome included a distinct gene (the ABC-transporter coding gene) (Fig. 1D ii). Similar locus is also present in *P. gingivalis* where the gene coding for the ABC transporter is followed by a gene coding for an outer membrane lipoprotein sorting protein (PG1179) and hypothetical protein (PG1178). An interesting finding was the PG1177 coding for a transposase followed by PG1176-75 encoding orthologs of BT0508-509 (Fig. 1D ii). A mobile element-encoding gene, BT1138 that has similar genes in the other two bacteria tested was also regulated in both studies. In addition, the BT3053 – 3055 encoding succinate dehydrogenase/fumarate reductase SdhABC was downregulated in both studies (Table 7, Fig. 2A). Finally, BT4715 *dps* coding for a DNA starvation/stationary phase protection protein is present in all three bacteria investigated in our study. It is of interest that in *B. thetaiotaomicron* upstream of *dps* is an *oxyR* (BT4716) facing in an opposite direction (Table 7) indicating that it may play a role in *dps* expression.

There were *B. thetaiotaomicron* loci that were regulated by iron levels and colitis and had counterparts in only one of the other bacteria investigated in our studies. The BT0503-504 locus has similar composition in *B. thetaiotaomicron* and *P. gingivalis* where the first gene coding for the hypothetical protein (BT0503) is the same in both bacteria and is followed by a TonB-encoding gene (BT0504 and PG2008), that although different in sequence, both code for functionally similarly TonB-dependent protein. Similarly, the BT0519-520 codes for a two-component hemerythrin-domain regulatory protein (Table 7, Fig. 3F). The operon is also present in *P. gingivalis* W83 and codes for the CdhR regulator ^25^. In addition, the BT0922-0923 coding for PepSY-like domain – containing protein as well as the BT2038-2040 encoding heavy metal efflux transporter system were present in both, *B. thataiotaomicron* and *P. gingivalis* (Table 7).

Finally, there were loci regulated by iron and colitis condition that are only present in *B. thetaiotaomicron* (Table 7). This includes the locus BT0523 – 0524 coding for a sugar transferase with its cognate response regulator, BT0532 encoding anthranilate synthase component I family protein, BT0995 encoding PAS domain-containing sensor histidine kinase, locus BT1042-1045 coding for a SusD/RagB transporter system, BT1597 coding for a HAMP domain-containing histidine kinase (followed by second component of the system and a gene coding for a cation efflux pump, BEX), BT1895 – 1896 encoding an outer membrane leucine-rich receptor system that is highly upregulated in colitis (Table 7).

BT2778, a sigma-70 family RNA Polymerase sigma factor, the multigene locus BT3625-3633 coding for a putative fimbrillin synthesis/transport locus, BT3969 encoding heavy metal transporter, BT4227 encoding MfaI fimbrillin protein, and BT4236 coding for a two-component response regulator, and BT4771 encoding RNA polymerase ECF type sigma factor were only present in *B. thetaiotaomicron*, (Table 7).

Some *B. thetaiotaomicron* loci that had orthologs of selected genes present in other bacteria studied. Thus, the BT2063 – 2065 locus was present mainly in *B. thetaiotaomicron* with the exception of an ortholog of the BT2065 coding for a TonB-dependent receptor also present in *P. intermedia.* BT2450 is a first gene in a locus coding for a hypothetical protein, pyrogenic exotoxin B (BT2451), and two hypothetical proteins (BT2452-3). *pdnA,* a gene similar to the BT2451, is present in *P. gingivalis* (PG1427) and codes for a periodontain light chain, as well as in *P. intermedia* (PIOMA14_I_1549) and encodes C10 family peptidase (Supplemental Fig. 3). Also, while BT4233 was only present in *B. thetaiotaomicron,* BT4234 had an ortholog in *P. intermedia*.

Finally, highly upregulated in both studies^2^ (iron levels and colitis) was a locus: BT2473 – BT2482 fimbrillin-family protein, two cytochrome c biogenesis proteins CcsA, porin, thiol oxidoreductase, peptidase M75, (helix-turn-helix transcriptional regulator, and three hypothetical proteins 69.4 – 323.7fold upregulation, Tables 5, 7, Fig. 3G).

There were downregulated pathways, both in our study and in the colitis study. The main downregulated locus was BT0552-3 coding for a glutamate synthase (*glt*). Also, downregulated in both studies was gene coding for asparagine synthase. In addition, nitrogen regulator encoded by BT0545 was significantly downregulated. While the above genes were only found in *B. thetaiotaomicron,* the downregulated operon coding for succinate dehydrogenase/fumarate reductase is present in all three bacteria studied (Table 7).

## Discussion

Our parallel study of three bacterial species belonging to the phylum of Bacteroidetes gives insight into the iron homeostasis in those organisms. During infection and associated with that inflammation iron is scarce and thus the conditions in the host resemble that of the iron limited conditions as reported in our study^26, 27, 28, 29^. Most *in vitro* studies are performed under iron sufficient conditions and thus our study provides more accurate depiction of the adaptation to host environment. We further verified the latter by comparing our results with results obtained from studies done *in vivo* using a colitis model ^2^. Although we previously reported the *P. gingivalis* – iron-dependent transcriptome this work further extends that study as well as compares it to other bacteria belonging to Bacteroidetes with the ultimate goal of having a consensus on iron-dependent regulation in this group of bacteria. We have previously reported *P. intermedia* proteins whose expression is affected by iron, however, this is the first comprehensive report on iron-dependent regulation in *P. intermedia* and the data reinforce our previously published findings ^16^ as well as data from other labs ^30^. The *B. thetaiotaomicron* iron-dependent regulation remains relatively unknown. Here we show comprehensive analysis of the stimulon using RNA derived from bacteria grown in iron replete and iron-chelated conditions. The number of regulated genes is larger than that found in *P. gingivalis* and *P. intermedia* that possibly is due to the much larger size of the genome of *B. thetaiotaomicron* ^31^ as well as the fact that *B. thetaiotaomicron* libraries were sequenced using the HiSeq Illumina sequencer while the other two library sets were sequenced using the MiSeq Illumina sequencer thus resulting in much larger number of reads for the former.

We do see changes in gene expression coding for proteins mediating several pathways: iron/heme acquisition, metabolism, and oxidative stress. Many of those pathways are shared among the three bacterial species. One of the most dramatically altered pathway is metabolism where the most drastically upregulated was a gene coding for flavodoxin, FldA. The FldA structure was first determined using the *E. coli* FldA ^32^, however, we note that the *P. gingivalis* FldA is more similar to *Disulfovibrio* FldA structure ^33^. Flavodoxins are electron transfer proteins that are present in variety of bacteria as well as algae and lower plants ^34, 35, 36^. Those are low molecular weight, FMN-containing proteins that function as electron transfer agents in a variety of microbial metabolic processes. Electron transfer proceeds through the exposed dimethylbenzene ring of the bound coenzyme FMN that is then reduced to FMNH_2_. FldA were isolated over a half century ago, first from Cyanobacteria and later from Clostridia^37, 38^ . FldA’s expression was shown to be induced under iron-limiting conditions, however, their expression can also be regulated by other stresses. Their biological role is similar to that of ferredoxins as FldA transfers electrons from donors to the electron transport chain components, starting with the Rnf complex, located in bacterial cell membrane thus creating sodium proton gradient driving ATP generation ^39^. While ferredoxins (Frds) use iron-sulfur clusters to transfer electrons, FldA binds the cofactor flavin mononucleotide (FMN) and thus is advantageous over ferredoxin as it spares iron for other essential enzymes relying on iron for their activity as well as is more resilient for oxidative/nitrosative stress damage. While we noted drastically upregulated FldA we also noted concomitant reduction in expression of ferredoxins, Frd under iron limiting conditions. Such iron-dependent expression was also noted in other organisms ^40, 34^. As inflammatory conditions lead to reduced iron levels as well as elevated oxidative stress mediators released by inflammatory cells, adaptation through elevation of FldA and replacement of Frd seems like an intuitive strategy benefiting persistence of the bacteria in such conditions. Our data may also have translational significance as FldA is highly upregulated in *B. thetaiotaomicron* in the colitis model as well as it’s transcript is very abundant in *P. gingivalis* identified in patients with periodontal disease ^24^. Such data point to a novel metabolic mechanism employed by *P. gingivalis* that relies more on iron independent enzymes during active disease status and thus contrasts with thus far favored metabolic model that is employed by the bacterium when grown in laboratory in iron rich conditions ^41^. It is noteworthy that FldA being an enzyme with characteristics unique to bacteria and not present in higher eukaryotes may also be a potential target for future development of therapeutic strategies ^36, 42^.

The other highly upregulated pathway includes an iron/heme acquisition mechanisms. Here, the well-defined system, the hemin uptake operon, *hmu* was observed to be upregulated in all three bacterial species tested. While in *P. gingivalis* and *B. thetaiotaomicron* one operon/genome was identified, the *P. intermedia* OMA14 displayed an *hmuY-*only carrying locus on the large chromosome (Ch I) and *hmu* operon on its small chromosome (Ch II), respectively (Table 3). Interestingly, in the *P. intermedia* 17 the *hmu* loci are also duplicated and present on separate chromosomes, but the orphan *hmuY* is located on the small chromosome (ChII) while the complete operon is present in the ChrI (Supplemental Fig. 1). The latter observation is consistent with the recent report from the Olczak laboratory ^30, 43^. We observed drastic upregulation of the complete *hmu* operon in all the three bacteria as well as the orphan *P. intermedia* 17 *hmuY* gene. Such findings are also in line with our previous report on iron-dependent proteome in *P. intermedia* 17 ^16^. It is noteworthy that this operon is also upregulated in *B. thetaiotaomicron* under colitis conditions when compared to healthy status of the animals ^2^. In periodontitis, hepcidin levels leading to reduction of systemic iron are elevated^44, 26^. Accordingly, in periodontitis, microbial iron uptake mechanisms are significantly upregulated when compared to healthy sites ^45, 46^. While iron acquisition through transport of hemin using the *hmu* system would seem like an intuitive strategy in low extracellular iron conditions we also observed upregulation of two putative transporter systems that include a TonB-dependent proteins and/or ABC-type transporters. The high upregulation in iron deficient conditions of those systems in most of the bacterial species tested warrants further investigation in their role in mediating growth and survival of the bacteria under low iron conditions. One such system BT2063-BT2065 has recently been characterized in *B. thetaiotaomicron* and shown to play a crucial role in iron acquisition through transport of xenosiderophores produced by *Salmonella typhimurium* ^2^. This locus is the most drastically upregulated in *B. thetaiotaomicron* and has no similarity (orthologs) in the other two bacteria studied. It is also noteworthy that this is the most upregulated locus in *B. thetaiotaomicron* derived from inflammatory condition (colitis model) when compared to non-inflamed host ^2^ thus further proving the biological significance of our findings.

The high upregulation of the loci with their composition including ABC type transporters and/or TonB dependent proteins also points to their role in possible iron acquisition from siderophores abundant in the oral or gut microbiomes. For the oral bacteria, such as *P. gingivalis* and *P. intermedia,* acquisition of iron from lactoferrin abundant in oral environment would seem like a probable strategy ^47, 3^. It would also be substantiated by the findings that bacterial siderophores have higher affinity for iron Fe(III) than lactoferrin ^48, 49^. It is thus probable that the ability to “steal” siderophores produced by other bacteria is present in both *P. gingivalis* and *P. intermedia* as those are bacteria residing in the complex oral microbiome where siderophore providers are abundant ^48^.

We found proteins within a multigene loci with similar functions across the three bacteria studied despite them bearing low identity at the amino acid level. Examples of that may be the CdhR-carrying locus. Furthermore, the order of genes within loci may have differed. An example is the locus PG1180-81 coding for a transporter and associated with it regulator that differ among the bacteria tested. The distribution of the genes within the locus differs as well and thus looking at pathways that are regulated in the context of a genome rather than searching for individual genes was a better prediction of the mechanisms that are deregulated during adaptation to low iron conditions.

Differences between the organisms were also detected and mainly included regulation of genes mediating carbohydrate utilization in *B. thetaiotaomicron.* This maybe due to the different niches in which the bacteria live; while *B. thetaiotaomicron* relies on carbohydrate utilization ^50^. Bacteroidetes have elaborate enzyme combinations that they use for glycan breakdown ^37, 51^*. P. gingivalis* and *P. intermedia* are asaccharolytic bacteria that have smaller genomes with 2.3Mb and 2.2 - 2.8 Mb (plus a mini genome ranging from 0.6 -0.9 Mb), respectively while the size of the genome of *Bacteroides thetaiotaomicron* is 6.26Mb (VPI-4582 strain) and thus is over twice as large^18, 52, 53, 31^. Both are lacking genes coding for glycan-degrading enzymes and thus *P. gingivalis* and *P. intermedia* rely on peptides for energy generation ^54, 18, 31^.

While all bacteria examined were able to reduce nitrite, that ability was iron- dependent in *B. thetaiotaomicron.* Furthermore, all bacteria tested have the nitrite reductase system, NrfAH, and only the *B. thetaiotaomicron* operon is drastically downregulated in iron limited conditions we thus we hypothesize that this operon plays a major role in physiology of that bacterium. Although reduction of expression and nitrite consumption was observed in the bacterium grown in the presence of iron chelator *B. thetaiotaomicron* is still able to consume large amounts of nitrite under iron- limiting conditions. Such capability may assist not only *B. thetaiotaomicron* but also *P. gingivalis* and *P. intermedia* to detoxify toxic byproducts of nitrate reduction to survive in the host. The nitrite levels in oral cavity have been shown to exceed 1mM, especially after nitrate-rich meal ^55, 56, 57^. Furthermore, one of the host’s defense mechanisms deployed by immune cells is production of nitric oxide that in turn can be oxidized to nitrite and thus effective nitrosative stress detoxification mechanisms would be expected in those bacteria.

We also detected large number of small genes that were regulated by iron. The role of those genes, coding for DUF1661 domain-containing proteins, is yet to be established. It is noteworthy that those genes are specific to *P. gingivalis* as no similarity to those genes in other bacteria was detected.

The striking similarity between genes upregulated under iron limited conditions in our study and the genes upregulated in colitis as oppose to healthy subjects points to iron being the master regulator of Bacteroides adaptation to the host inflammatory condition. So far the knowledge of the molecular mechanisms of regulating expression of the effector proteins required for adaptation of Bacteroidetes to differing iron levels remain scarce. The identification of the above commonly regulated genes in all the three bacterial species is predicted to help identify iron-dependent regulatory systems in those bacteria. CdhR has been shown to regulate expression of *hmuYR* ^25, 58^, however, the molecular mechanisms of the regulation remain unclear as CdhR has yet to be shown to bind iron. Furthermore, given the clinical correlation between genes expressed in iron limited conditions and genes highly expressed during active inflammatory disease, the mechanisms identified in our study to be drastically upregulated gain biological significance what warrants further detailed investigation of their function and molecular mechanisms.

In conclusion, we have compared the iron-dependent stimulons of three bacterial species belonging to the *Bacteroidetes* phylum. In addition to the iron update mechanisms, the bacteria exhibited major adaptation of their metabolic mechanisms as well as oxidative stress response mechanisms. The iron-limited conditions resembled those found in inflamed host and thus indicate that iron is the major player in regulation of bacterial adaptation to diseased conditions. Such finding warrants more in depth investigation of the molecular mechanisms of the iron-based effector proteins as well as the regulatory mechanisms that thus far remain elusive in the Bacteroidetes phylum.

### Data Availability

The RNAseq data has been deposited to the NCBI Gene Expression Omnibus (GEO). The data are available under the accession numbers: GSE169260 (for *B. thetaiotaomicron* BT5482 Δ*tdk*), GSE168982 (for *P. intermedia* OMA14), GSE168570 (for *P. gingivalis* W83), and GSE174493 (for *P. gingivalis* ATCC33277).

## Acknowledgements

This work was funded by NIH grants 1R01DE023304 (J.P.L.), NIHR21DE025555 (J.P.L.) and K18 DE029865 (J.P.L.). Figures were generated using the BioCyc Pathways Tools version 26.0 (software by SRI International).

## Author Contributions

J.P.L. performed experiments and analyzed all data. J.P.L. wrote the manuscript. Q.G. performed experiments.

## Supplementary Figures

Supplemental Figure 1. Iron-regulated loci in *Prevotella intermedia* OMA14. Iron-regulated loci in *Prevotella intermedia* OMA14. A. Loci upregulated in low iron. *i. hmu* – full operon*. ii. fldA* – locus (PIOMA14_I_0020). iii. RS06100 *hmuY* (PIOMA14_I_1196)*–* one gene operon comparison with the same locus in *P. intermedia* 17 and *P. gingivalis* W83. B. Loci downregulated in low iron. i. *sdh* – locus (located on chromosome I), PIOMA14_II_0616, ii. RS13160-13200 *rnf* electron transport – PIOMA14_I_0607 – 0616, iii. PS03360-80 *vor* locus -PIOMA_I_0666 – 0670. iv. RS10750 *rbr* locus – PIOMA14_II_0105. v. RS05315 *frd* (4Fe-4S binding protein) – PIOMA14_I_1049.

Supplemental Figure 2. Other loci of interest in *P. intermedia* OMA14. i. C10-family peptidase coding genes. ii. *nrfAH* locus. PIOMA14_I_0579 – 0582. RS00350 – 355 *ahpCF* locus – PIOMA14_I_0069 – 70. RS09985 *dps* locus PIOMA14_I_1962.

Supplemental Figure 3. Other interesting loci in Bacteroidetes. i. toxin/peptidase encoding gene. ii. *dps* coding for the DNA starvation/stationary phase protection protein.

Supplemental Figure 4. Small iron-regulated proteins in Bacteroidetes. *P. gingivalis* W83 DUF1661 domain-containing proteins.

## Supplementary Tables

Table 1. Genes upregulated in P. gingivalis ATCC33277 at least 2 fold under iron deplete conditions.

Table 2. Genes downregulated in P. gingivalis ATCC33277 at least 2 fold under iron deplete conditions.

